# Improved radiation expression profiling in blood by sequential application of sensitive and specific gene signatures

**DOI:** 10.1101/2021.08.18.456812

**Authors:** Eliseos J. Mucaki, Ben C. Shirley, Peter K. Rogan

## Abstract

**Purpose:** Combinations of expressed genes can discriminate radiation-exposed from normal control blood samples by machine learning based signatures (with 8 to 20% misclassification rates). These signatures can quantify therapeutically-relevant as well as accidental radiation exposures. The prodromal symptoms of Acute Radiation Syndrome (ARS) overlap those present in Influenza and Dengue Fever infections. Surprisingly, these human radiation signatures misclassified gene expression profiles of virally infected samples as false positive exposures. The present study investigates these and other confounders, and then mitigates their impact on signature accuracy.

**Methods:** This study investigated recall by previous and novel radiation signatures independently derived from multiple Gene Expression Omnibus datasets on common and rare non-malignant blood disorders and blood-borne infections (thromboembolism, S. aureus bacteremia, malaria, sickle cell disease, polycythemia vera, and aplastic anemia). Normalized expression levels of signature genes are used as input to machine learning-based classifiers to predict radiation exposure in other hematological conditions.

**Results:** Except for aplastic anemia, these blood-borne disorders modify the normal baseline expression values of genes present in radiation signatures, leading to false-positive misclassification of radiation exposures in 8 to 54% of individuals. Shared changes, predominantly in DNA damage response and apoptosis-related gene transcripts in radiation and confounding hematological conditions, compromise the utility of these signatures for radiation assessment. These confounding conditions (sickle cell disease, thromboembolism, S. aureus bacteremia, malaria) induce neutrophil extracellular traps, initiated by chromatin decondensation, DNA damage response and fragmentation followed by programmed cell death. Riboviral infections (for example, Influenza or Dengue fever) have been proposed to bind and deplete host RNA binding proteins, inducing R-loops in chromatin. R-loops that collide with incoming replication forks can result in incompletely repaired DNA damage, inducing apoptosis and releasing mature virus. To mitigate the effects of confounders, we evaluated predicted radiation-positive samples with novel gene expression signatures derived from radiation-responsive transcripts encoding secreted blood plasma proteins whose expression levels are unperturbed by these conditions.

**Conclusions:** This approach identifies and eliminates misclassified samples with underlying hematological or infectious conditions, leaving only samples with true radiation exposures. Diagnostic accuracy is significantly improved by selecting genes that maximize both sensitivity and specificity in the appropriate tissue using combinations of the best signatures for each of these classes of signatures.

## Introduction

One of the most promising approaches to quantify absorbed ionizing radiation is based on levels of different gene expression responses in blood (Dressman et al. 2007; Paul and Amundson, 2008; Ding et al. 2013; Lu et al. 2014). Combinations of gene expression levels, termed signatures, can predict ionizing radiation exposure in humans and mice from publicly available microarray gene expression levels (Zhao et al. 2018a). Several groups (Boldt et al. 2012; Budworth et al. 2012; Knops et al. 2012; Ghandhi et al. 2017) have also used signatures to determine radiation exposures. Our approach uses supervised machine learning (ML) with genes previously implicated or established from genetic evidence and biochemical pathways that are altered in response to these exposures (Zhao et al. 2018a). Biochemically-inspired ML is a robust approach to derive diagnostic gene signatures for radiation and chemotherapy (Dorman et al. 2016; Mucaki et al. 2016; Mucaki et al. 2019; Bagchee-Clark et al. 2020). Given the limited sample sizes of typical datasets, appropriate ML methods for deriving gene signatures have included Support Vector Machines, Random Forest classifiers, Decision Trees, Simulated Annealing, and Artificial Neural Networks (Rogan, 2019; Boldrini et al. 2019).

Before performing ML, we ranked a set of curated radiation-response genes in radiation- exposed samples by Minimum Redundancy Maximum Relevance (mRMR; Ding and Peng; 2005), which orders genes based on mutual information between their expression and whether the sample was irradiated (MR) and the degree to which their expression profile is dissimilar from previously selected genes (mR). A Support Vector Model (SVM) or signature is generated from the top ranked gene set by adding and removing genes that minimize either model misclassification or log-loss (Zhao et al. 2018a). ML alone is not sufficient to derive useful signatures, since underlying biochemical pathways and disease mechanisms also contribute to selecting relevant and non-redundant gene features (Bagchee-Clark et al. 2020). Signatures were evaluated either by a traditional validation approach or by stratified k-fold validation (which splits the same dataset into k groups reserved for testing and training). The traditional model-centric approach uses a normalized training set which is then used to predict outcome of normalized independent test data. While more susceptible to batch effects than k-fold validation, this approach can incorporate smaller datasets or heterogeneous data sources.

The present study is concerned with the selection of "normal" controls for training and testing signatures. Selection of appropriate controls has been a debated subject in other medical fields (Lipsitch et al. 2011). In individuals with clinically overlapping diagnoses, this has a critical but underappreciated impact on the accuracy of molecular diagnostic testing. Previous studies have revealed gene expression changes in blood from unirradiated individuals with underlying metabolic or confounders, including smokers (Paul and Amundson, 2011) and modulators of inflammation such as lipopolysaccharide of bacterial origin or curcumin (Cruz-Garcia et al. 2018). There is also evidence of interactions between hematological comorbidities and radiation exposure. Radiodermatitis has been associated with S. aureus infections (Hill et al. 2004). Radiation therapy has also been contraindicated in individuals with venous thromboembolism (Guy et al. 2017).

While investigating the possibility of using radiation gene signatures to differentiate the prodromal phase of Acute Radiation Syndrome (ARS) from early-stage Influenza, riboviral infections induced expression changes similar to those seen in gamma-irradiated samples (Rogan et al. 2021). Radiation signatures misclassified some unirradiated blood samples from infected individuals as radiation-exposed (derived in Zhao et al. 2018a; designated M1-M4 in Rogan et al. 2021). False positive radiation exposure predictions of unirradiated samples from individuals diagnosed of Influenza has also been noted by others (Jacobs et al. 2020 [Supplementary data]). The M1-M4 signatures consisted predominantly of genes with roles in DNA damage response, programmed cell death, and inflammation. However, many of these genes were also similarly dysregulated in blood from Influenza and Dengue virus-infected samples. The expression of the DNA damage response gene *DDB2*, for example, increases with radiation, but was also induced in a significant number of viral infected samples which were then classified incorrectly as radiation exposed (Figure 5 of Rogan et al. 2021). *DDB2* is present in many other radiation gene signatures or developed radiation-exposure assays (Paul and Amundson, 2008; Lu et al. 2014; Jacobs et al. 2020).

While this process strictly validates signatures to estimate sensitivity to radiation, the same rigor has not been applied to determining specificity, which we suspected could be impacted by comorbidities in the general population. This study considers other hematological conditions that alter the normal expression of the same genes in blood that are often selected for assessment of radiation exposure and suggests an approach for addressing this issue.

We evaluated gene expression data from other individuals with blood-borne conditions (infections, inherited and idiopathic hematological disorders) to determine whether previous and novel gene signatures could discriminate radiation exposure from these phenotypes. We investigate whether these effects are reproducible using newly derived signatures from independent datasets derived from irradiated blood samples (Figure 1).

**Figure 1:**
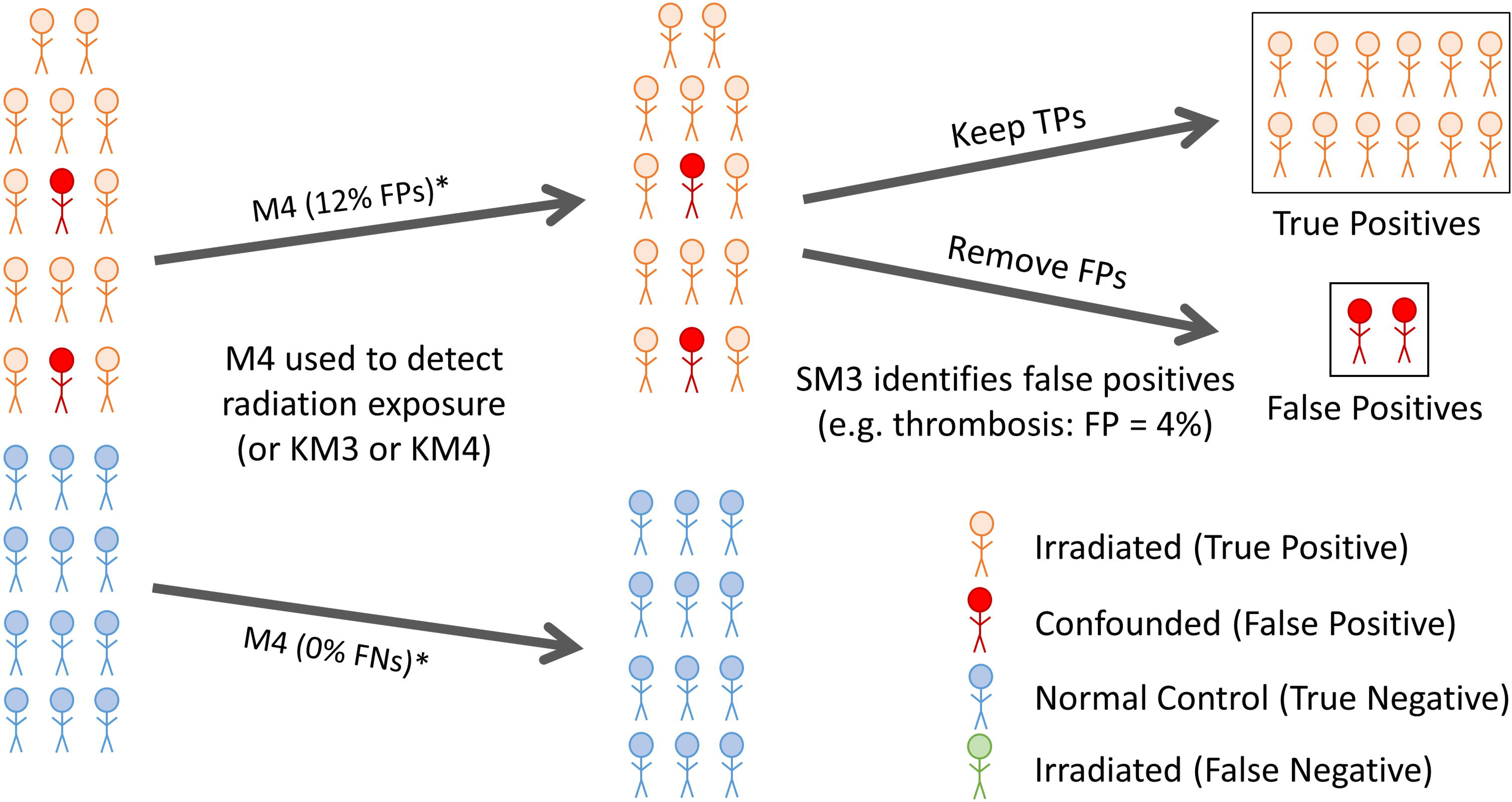
Evaluation of Conditions Confounding Radiation Gene Signatures. The traditional validation approach was used to evaluate unirradiated datasets for hematological conditions to assess the performance of radiation gene signatures derived in Zhao et al. (2018a). With this approach, we can identify and then reject models with high rates of FP radiation diagnosis in confounding conditions while identifying what confounders could make individuals ineligible for a radiation gene signature assay. Alternatively, new radiation gene signatures could be derived that show improved FP rates in both controls and test subjects.

## Methods

### Datasets evaluated

Expression data from the Gene Expression Omnibus [GEO; https://www.ncbi.nlm.nih.gov/geo/] and Array Express [https://www.ebi.ac.uk/arrayexpress/] databases were required to contain the same genes in each signature in both training and independent validation radiation datasets (Zhao et al. 2018a). These datasets were: GSE1725 [Designated: RadLymphCL-1], GSE6874 [GPL4782; Designated: RadTBI-2], GSE10640 [GPL6522; Designated: RadTBI-3] and GSE701 [Designated: RadLymphCL-4] (Table 1). Several well known radiation genes which have appeared in other radiation gene signatures (Paul and Amundson, 2008; Oh et al. 2014; Port et al. 2017; Tichy et al. 2018; Jacobs et al. 2020) were previously not considered and were not present in any of our former ML models. Genes were excluded either because they were: 1) absent from one or more datasets (e.g. *FDXR*, *RPS27L*, *AEN* were missing from RadTBI-3); 2) mislabeled in the dataset with a legacy name leading to a mismatch between datasets (e.g. *PARP1* appears as *ADPRT* in RadTBI- 2); 3) secondary RNAs, such as micro- or long non-coding-RNA derived from the same gene (e.g. *BBC3* probes also detected multiple microRNAs in RadLymphCL-1; *POU2AF1* probes in RadLymphCL-4 indicated as LOC101928620’); or 4) missing from the set of curated radiation response genes (e.g. *PHPT1*, *VWCE*, *WNT3*).

**Table 1.**
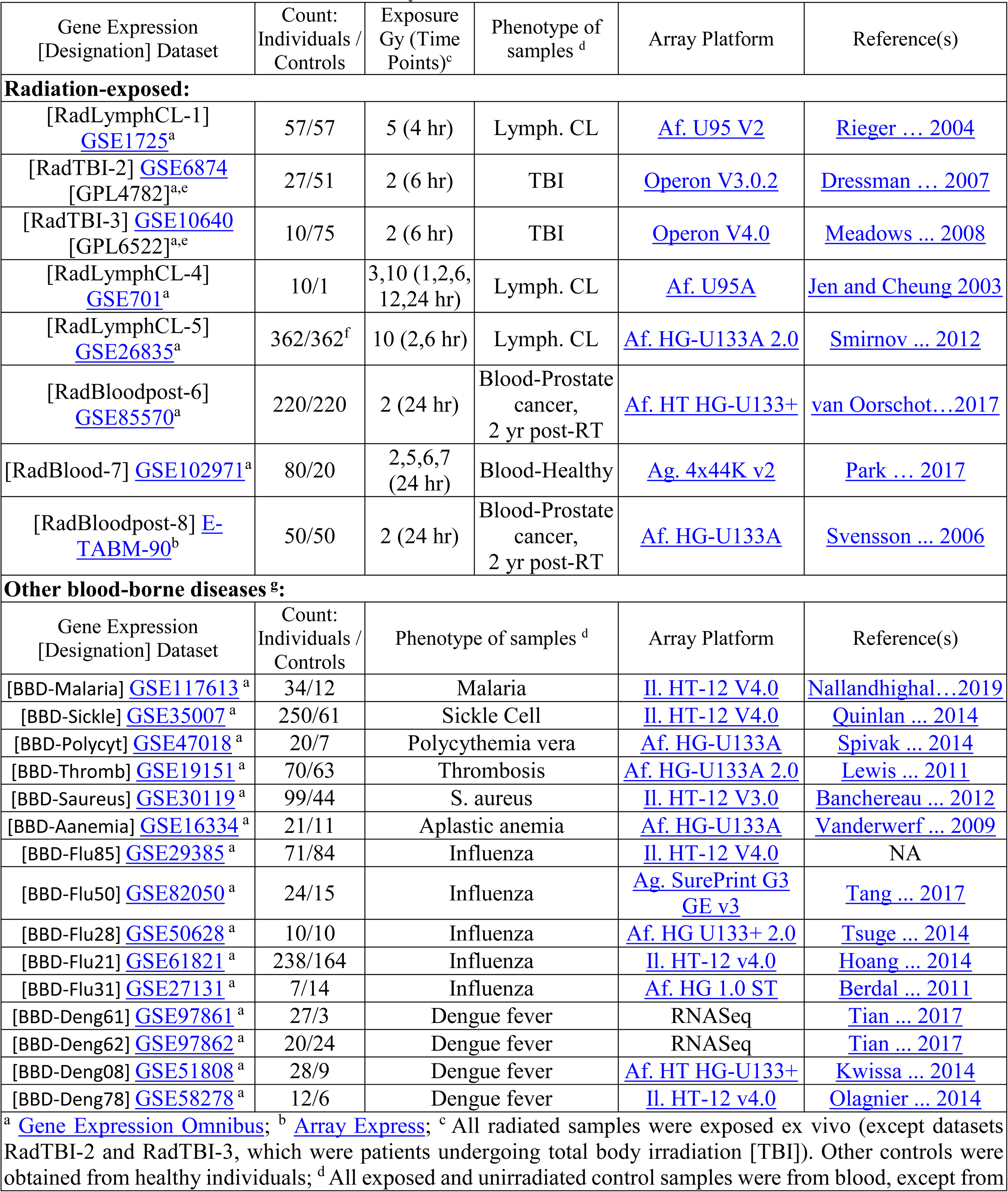
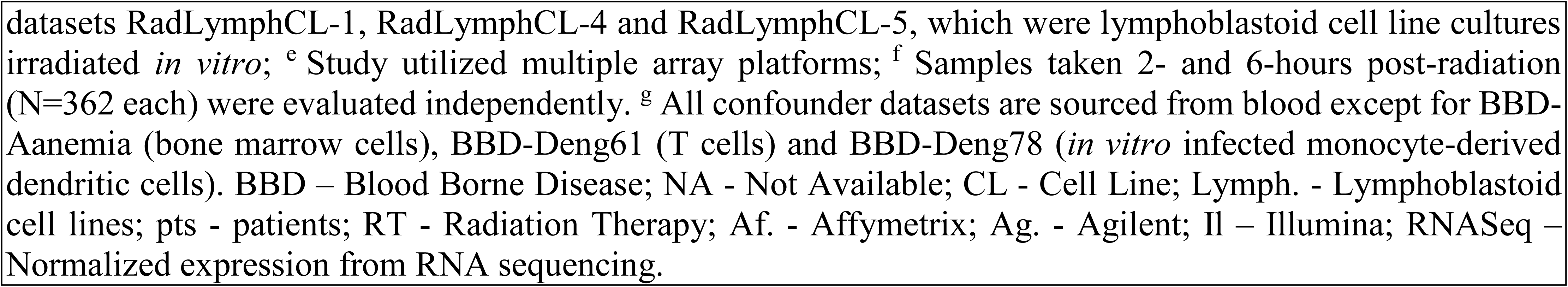
Characteristics of Datasets Analyzed.

| Gene Expression<br>[Designation] Dataset | Count:<br>Individuals /<br>Controls | Exposure<br>Gy (Time<br>Points)<sup>c</sup> | Phenotype of<br>samples <sup>d</sup> | Array Platform | Reference(s) |
| --- | --- | --- | --- | --- | --- |
| **Radiation-exposed:** |  |  |  |  |  |
| [RadLymphCL-1]<br>GSE1725<sup>a</sup> | 57/57 | 5 (4 hr) | Lymph. CL | Af. U95 V2 | Rieger ... 2004 |
| [RadTBI-2] GSE6874<br>[GPL4782]<sup>a,e</sup> | 27/51 | 2 (6 hr) | TBI | Operon V3.0.2 | Dressman ... 2007 |
| [RadTBI-3] GSE10640<br>[GPL6522]<sup>a,e</sup> | 10/75 | 2 (6 hr) | TBI | Operon V4.0 | Meadows ... 2008 |
| [RadLymphCL-4]<br>GSE701<sup>a</sup> | 10/1 | 3,10 (1,2,6,<br>12,24 hr) | Lymph. CL | Af. U95A | Jen and Cheung 2003 |
| [RadLymphCL-5]<br>GSE26835<sup>a</sup> | 362/362<sup>f</sup> | 10 (2,6 hr) | Lymph. CL | Af. HG-U133A 2.0 | Smirnov ... 2012 |
| [RadBloodpost-6]<br>GSE85570<sup>a</sup> | 220/220 | 2 (24 hr) | Blood-Prostate<br>cancer,<br>2 yr post-RT | Af. HT HG-U133+ | van Oorschot...2017 |
| [RadBlood-7] GSE102971<sup>a</sup> | 80/20 | 2,5,6,7<br>(24 hr) | Blood-Healthy | Ag. 4x44K v2 | Park ... 2017 |
| [RadBloodpost-8] E-TABM-90<sup>b</sup> | 50/50 | 2 (24 hr) | Blood-Prostate<br>cancer,<br>2 yr post-RT | Af. HG-U133A | Svensson ... 2006 |
| **Other blood-borne diseases <sup>g</sup>:** |  |  |  |  |  |
| Gene Expression<br>[Designation] Dataset | Count:<br>Individuals /<br>Controls | Phenotype of samples <sup>d</sup> |  | Array Platform | Reference(s) |
| [BBD-Malaria] GSE117613<sup>a</sup> | 34/12 | Malaria |  | II. HT-12 V4.0 | Nallandhighal...2019 |
| [BBD-Sickle] GSE35007<sup>a</sup> | 250/61 | Sickle Cell |  | II. HT-12 V4.0 | Quinlan ... 2014 |
| [BBD-Polycyt] GSE47018<sup>a</sup> | 20/7 | Polycythemia vera |  | Af. HG-U133A | Spivak ... 2014 |
| [BBD-Thromb] GSE19151<sup>a</sup> | 70/63 | Thrombosis |  | Af. HG-U133A 2.0 | Lewis ... 2011 |
| [BBD-Saureus] GSE30119<sup>a</sup> | 99/44 | S. aureus |  | II. HT-12 V3.0 | Banchereau ... 2012 |
| [BBD-Aanemia] GSE16334<sup>a</sup> | 21/11 | Aplastic anemia |  | Af. HG-U133A | Vanderwerf ... 2009 |
| [BBD-Flu85] GSE29385<sup>a</sup> | 71/84 | Influenza |  | II. HT-12 V4.0 | NA |
| [BBD-Flu50] GSE82050<sup>a</sup> | 24/15 | Influenza |  | Ag. SurePrint G3<br>GE v3 | Tang ... 2017 |
| [BBD-Flu28] GSE50628<sup>a</sup> | 10/10 | Influenza |  | Af. HG U133+ 2.0 | Tsuge ... 2014 |
| [BBD-Flu21] GSE61821<sup>a</sup> | 238/164 | Influenza |  | II. HT-12 v4.0 | Hoang ... 2014 |
| [BBD-Flu31] GSE27131<sup>a</sup> | 7/14 | Influenza |  | Af. HG 1.0 ST | Berdal ... 2011 |
| [BBD-Deng61] GSE97861<sup>a</sup> | 27/3 | Dengue fever |  | RNASeq | Tian ... 2017 |
| [BBD-Deng62] GSE97862<sup>a</sup> | 20/24 | Dengue fever |  | RNASeq | Tian ... 2017 |
| [BBD-Deng08] GSE51808<sup>a</sup> | 28/9 | Dengue fever |  | Af. HT HG-U133+ | Kwissa ... 2014 |
| [BBD-Deng78] GSE58278<sup>a</sup> | 12/6 | Dengue fever |  | II. HT-12 v4.0 | Olagnier ... 2014 |
| <sup>a</sup> Gene Expression Omnibus; <sup>b</sup> Array Express; <sup>c</sup> All radiated samples were exposed ex vivo (except datasets RadTBI-2 and RadTBI-3, which were patients undergoing total body irradiation [TBI]). Other controls were obtained from healthy individuals; <sup>d</sup> All exposed and unirradiated control samples were from blood, except from |  |  |  |  |  |

datasets RadLymphCL-1, RadLymphCL-4 and RadLymphCL-5, which were lymphoblastoid cell line cultures irradiated *in vitro*; <sup>e</sup> Study utilized multiple array platforms; <sup>f</sup> Samples taken 2- and 6-hours post-radiation (N=362 each) were evaluated independently. <sup>g</sup> All confounder datasets are sourced from blood except for BBD-Aanemia (bone marrow cells), BBD-Deng61 (T cells) and BBD-Deng78 (*in vitro* infected monocyte-derived dendritic cells). BBD – Blood Borne Disease; NA - Not Available; CL - Cell Line; Lymph. - Lymphoblastoid cell lines; pts - patients; RT - Radiation Therapy; Af. - Affymetrix; Ag. - Agilent; Il – Illumina; RNASeq – Normalized expression from RNA sequencing.

To address the possibility that inclusion of genes missing from signatures based on RadLymphCL-1, RadTBI-2 or RadTBI-3 might improve radiation response prediction, we developed novel signatures from more recent radiation datasets, including GEO: GSE26835, GSE85570, GSE102971 and ArrayExpress: E-TABM-90 (Designated RadLymphCL-5, RadBloodpost-6, RadBlood-7, and RadBloodpost-8, respectively; Table 1). Previous radiation gene signatures (Zhao et al. 2018a) were derived from GSE6874[GPL4782] and GSE10640[GPL6522] (RadTBI-2 and RadTBI-3, respectively; bracketed accession numbers refer to the subset of samples belonging to a corresponding GEO SuperSeries). Exposure levels were at a minimum of 2Gy (including total body irradiation [TBI]) and were carried out between 4-24 hours post-irradiation for all radiation-derived signatures.

To investigate whether other disorders could also confound radiation signatures, we assessed performance of radiation signatures utilized in this study with available gene expression datasets for other blood-borne diseases. These datasets include GEO: GSE117613 (cerebral <u>malaria</u> and severe malarial anemia; Designated: BBD-Malaria [BBD – Blood Borne Disease]), GSE35007 (<u>sickle cell disease</u> in children; Designated: BBD-Sickle), GSE47018 (polycythemia vera; Designated: BBD-Polycyt), GSE19151 (single and recurrent venous <u>thromboembolism</u>; Designated: BBD-Thromb), GSE30119 (Staphylococcus [<u>S.</u>] <u>aureus</u> infection; Designated: BBD-Saureus), and GSE16334 (aplastic anemia; Designated: BBD-Aanemia) (Table 1). Gene expression was measured by microarray in each dataset (qRT-PCR validation data was not available). An idiopathic portal hypertension dataset (GSE69601) was excluded due to insufficient numbers of samples (N=6).

### Data Preprocessing

Microarray data from each dataset were pre-processed as described in Zhao et al. (2018a). Briefly, missing gene expression values were imputed (gene feature was removed if values were missing from >5% samples) from nearest neighbours, the expression values of patient replicates were averaged, and gene expression of all genes were z-score normalized. Genes previously implicated in the radiation response (N=998) were analyzed, including 13 additional radiation genes described in other studies, including *CD177*, *DAGLA*, *HIST1H2BD*, *MAMDC4*, *PHPT1*, *PLA2G16*, *PRF1*, *SLC4A11*, *STAT4*, *VWCE*, *WLS*, *WNT3*, and *ZNF541* (N=1,011 genes total).

### Derivation of Radiation Gene Signatures

mRMR ranking was determined for curated, expressed radiation-related genes in the presence or absence of radiation. mRMR first selects the gene with the highest mutual information (MI; Cover and Thomas, 2006; Zeng 2015) between its expression level and the radiation exposure status of each sample. MI ranges from 0 to 1 bit for a gene for a set comprised of radiated and unirradiated samples and measures the mutual dependence between the radiation exposure status and expression for each gene within the same dataset. With MI = 1 bit, expression levels and radiation exposure are perfectly correlated. Expression levels of a gene that distinguish some but not all irradiated and unirradiated samples produce MI values between 0 and 1. A low MI value ∼0 bits indicates that expression levels are weakly or uncorrelated with the radiation phenotype. mRMR feature selection then minimizes redundant expression patterns among the genes chosen by prioritizing gene candidates with the highest difference between its MI and the average MI of all previously selected genes with the candidate gene as a probability vector (Ding and Peng, 2005; Mucaki et al. 2016). Minimizing redundancy results in some subsequent selected gene(s) with orthologous expression patterns relative to the preceding gene(s). These may exhibit significantly lower MI, typical of a weak radiation response. Nevertheless, higher ranked gene features exhibit larger MI values in general. Gene rankings by mRMR and the computed MI for each radiation gene in each of the datasets evaluated are provided in Suppl. Table S1.

SVM-based gene signatures were derived by greedy feature selection, including forward sequential feature selection (FSFS), backward sequential feature selection (BSFS) and complete sequential feature selection (CSFS; Zhao et al. 2018a). Our software for biochemically-inspired ML is available in a Zenodo archive (Zhao et al. 2018b). Both FSFS and BSFS models were derived from the top 50 ranked mRMR genes, in addition to other published radiation responsive genes: *AEN*, *BAX*, *BCL2*, *DDB2*, *FDXR*, *PCNA*, *POU2AF1*, and *WNT3*. SVMs were derived with a Gaussian radial basis function kernel by iterating over box-constraint (C) and kernel-scale (*σ*) parameters and gene features, minimizing to either misclassification or log loss by cross-validation (Zhao et al. 2018a; Bagchee-Clark et al. 2020). Gene signatures were then assessed with a validation dataset and re-evaluated (by misclassification rates, log loss, Matthews correlation coefficient, or goodness of fit). This study primarily reports misclassification rates to simplify comparisons of results between radiation-exposed and disease confounder datasets. Those with high misclassification rates in validated radiation datasets (>50%) have been excluded. Variation in radiation signature composition among different source datasets can be attributed to distinct microarray platforms, other batch effects, and inter-individual variation in gene expression which cannot be fully mitigated by normalization. These contribute to MI variability, which both alters mRMR rank and features selected during signature derivation.

The quality of each dataset was assessed based on the dynamic range in responses by signature genes to radiation. This was based on the premise that these responses among different confounding datasets might alter expression of some of the same radiation-responsive genes. MI between gene expression and radiation dose is indicated in Suppl. Table S1. Datasets RadBloodpost-6 and RadBlood-7 both exhibit high MI with radiation exposure (maximum MI > 0.7 bits for both datasets; 77 and 115 genes with > 0.2 bits MI, respectively). By contrast, datasets RadLymphCL-5 and RadBloodpost-8 both exhibited low MI for top ranked genes with radiation exposure. Of the top 50 ranked genes in dataset RadBloodpost-8, 13 genes had MI values <10% of the MI of the top ranked gene [0.3 bits], and 856 of 860 genes in the complete dataset had MI < 0.2 bits. The maximum MI for RadLymphCL-5 was 0.25 bits; 40 of 50 top ranked genes exhibited <10% MI of this value, and the radiation response genes *DDB2*, *PCNA*, *FDXR*, *AEN*, and *BAX,* had unexpectedly low rankings (>100; 2h and 6h post-exposure) and MI < 0.15 bits. The low MIs across all eligible genes indicates that the response to radiation was nearly random in both datasets. These datasets failed to satisfy minimum quality criteria and were excluded from further analyses. Radiation toxicity in dataset RadBloodpost-8 and cell line immortalization in RadLymphCL-5 appears to compromise their radiation response.

### Radiation gene signatures derived from expressed genes encoding secreted factors originating from blood cells

Only genes that encoded proteins present in blood plasma were used to derive an alternate set of gene expression signatures. The initial set of plasma proteins from the Human Protein Atlas “Human Secretome” [http://www.proteinatlas.org/humanproteome/secretome] and the Plasma Protein Database [http://www.plasmaproteomedatabase.org] were cross-referenced to create a list of 1377 shared proteins. The Genotype-Tissue Expression (GTEx) Portal (https://gtexportal.org/home/) was used to determine which genes encoding these secreted proteins are expressed in either leukocytes or transformed lymphoblasts at detectable levels (where Transcripts Per Million (TPM) > 1; N=682). Expressed genes present in radiation datasets RadTBI-2 (N=428) and RadTBI-3 (N=325) were used to derive ML models of genes encoding for human plasma proteins using CSFS, BSFS, and FSFS.

### Evaluating specificity of radiation gene signatures with expression of genes in confounding hematological conditions

Radiation gene signatures derived in this study (M5-M20) and in Zhao et al. (2018a) (M1-M7, KM3-KM7) were used to evaluate datasets consisting of independent expression measurements of the same gene features in samples derived from hematological disorders and controls (Table 1). Traditional ML validation was performed using available software (‘*regularValidation_multiclassSVM.m*’; Zhao et al. 2018b). This software performs quantile normalization to the features of the training and test set together (making the distributions of the two datasets statistically identical) before fitting a model to the training data which is then used to predict exposure based on the normalized expression of the test set. We evaluated how often these unirradiated individuals were misclassified as radiation-exposed by these models. Significant differences between false positive (FP) blood-borne cases and controls for the same signature were determined with the Mantel-Haenszel chi square and Mid-P exact tests, using a threshold of p = 0.05. Assessing unirradiated expression datasets featuring patients with hematological or infectious conditions using radiation signatures will not yield any true positive (TP) or false negative (FN) cases. This is because our experimental design did not merge radiation and confounder datasets by joint normalization of expression values. Normalization often does not adequately account for variations due to batch effects or between different microarray platforms. Furthermore, we cannot exclude that irradiated samples from healthy individuals may mask underlying blood-based phenotypes that were not documented in published studies. For these reasons, radiation and hematological disorder datasets were evaluated separately with the same signatures for FP and TN levels only, rather than by positive or negative predictive values. We determined the impact of genes in each signature on misclassification by iteratively removing individual gene features, rederiving signatures with these genes, and redetermining misclassification rates using expression data from hematological or infectious diseases (Zhao et al. 2018a; Mucaki et al. 2019). Expression levels of radiation responsive genes in confounder datasets of correctly vs. misclassified samples were contrasted using violin plots. These display weighted distributions of the normalized gene expression from each confounder datasets which were either properly (true negative [TN]) and improperly (FP) classified as irradiated by the radiation gene signatures (created in R [i386 v4.0.3] with *ggplot2*). Counts of confounder sub- phenotypes were stratified using Sankey diagrams (SankeyMATIC; http://sankeymatic.com/build/) which display the distribution of misclassified samples by phenotype. This analysis delineates FP and TN predictions (at the individual level) of groups of diseased patients and controls from these datasets according to predictions of the designated, specific radiation gene signature.

## Results

### Initial Evaluation of Candidate Genes in Radiation Gene Expression Datasets for Machine Learning

We derived new gene expression signatures by leave-one-out and k-fold cross-validation from microarray data based on more recent comprehensive gene datasets (RadBloodpost-6 and RadBlood-7) besides those we previously reported (Zhao et al. 2018a). Only some of the 1,011 curated genes were present on these microarray platforms, including 864 genes in RadBloodpost- 6 and 971 genes of RadBlood-7. After normalization, gene rankings by mRMR between datasets RadBloodpost-6 and RadBlood-7 were similar (Suppl. Table S1). In RadBloodpost-6, *FDXR* were ranked first, while *AEN* was top ranked in RadBlood-7 (*FDXR* was ranked 38th). *DDB2* was top ranked in datasets RadTBI-2 and RadTBI-3 (Zhao et al. 2018a), datasets which lacked expression for *FDXR* and *AEN,* respectively. Radiation-response genes among the top 50 ranked genes present in all 4 datasets included *BAX*, *CCNG1*, *CDKN1A, DDB2*, *GADD45A, PPM1D* and *TRIM22*.

*ERCC1* was selected by mRMR to be the second-ranked gene in dataset RadBlood-7, even though its MI was 31-fold lower than the top ranked gene, *AEN* (Suppl. Table S1). MI of the second-ranked genes in dataset RadTBI-2 (*RAD17*) was 7-fold lower than the first (*DDB2*), while dataset RadTBI-3 (*CD8A*) showed a 4-fold difference. Six of the top 50 genes in RadBlood-7 exhibited <10% of the MI of *AEN* (3 genes for RadTBI-2; none of the top 50 in RadTBI-3 and RadBloodpost-6 were <10% of the top ranked gene). Selection of low MI genes by ML feature selection likely reduces accuracy of gene signatures during validation steps. In the future, signature derivation will set a minimum MI threshold for ranking genes by mRMR.

The overall levels of MI for top ranked genes in datasets RadBloodpost-6 (0.72 bits for *AEN*) and RadBlood-7 (0.82 bits for *FDXR*) were comparable (Suppl. Table S1). In RadBlood-7, the genes with the highest MI were *AEN*, *DDB2*, *FDXR*, *PCNA* and *TNFRSF10B* (closely followed by *BAX*). While each were found in the top 50 ranked genes, some rankings were decreased to minimize redundant information (*FDXR* and *AEN* are ranked #38 and #41 in the RadBlood-7 dataset, respectively). MI for the top ranked genes in datasets RadTBI-2 and RadTBI- 3 were lower by comparison (0.31 and 0.47 bits for *DDB2*, respectively); the depressed maximum MI values in these datasets may, in part, be related to reduced numbers of eligible genes on these microarray platforms.

### Radiation Gene Signature Performance in Blood-Borne Diseases

The specificity of previously-derived radiation signatures selected after k-fold validation (KM1- KM7) and traditional validation (M1-M4; Zhao et al. 2018a) was assessed with normalized expression data of patients with unrelated hematological conditions, rather than evaluating unirradiated healthy controls. Signatures M1 and M2 (derived from dataset RadTBI-3; Table 2) and M3 and M4 (derived from RadTBI-2; Table 2) were assessed with multiple expression datasets from Influenza A (BBD-Flu##) and Dengue fever (BBD-Deng##) blood infections (Rogan et al. 2021; Table 1). FPs for radiation exposure were defined as instances where the misclassification rates of individuals with the disease diagnosis exceeded normal controls. A clear bias towards FP predictions of infected samples relative to controls was evident with all of these radiation gene signatures (Rogan et al. 2020; extended data – Section 1 Table 7). Dissection of the ML features responsible implicated 10 genes contributing to misclassification, including *BCL2*, *DDB2* and *PCNA*. These other conditions also confound predictions by radiation signatures derived by k-fold validation (KM1-KM7; Table 2).

**Table 2.** Traditional and K-Fold Validated Radiation Gene Expression Signatures M1-M4 and KM1-KM7.

| Signature |  | FS. Algorithm | Accuracy<sup>1</sup> |
| --- | --- | --- | --- |
| **a) Derived from Radiation Dataset RadLymphCL-1 (GSE1725)** |  |  |  |
| KM1 | *GADD45A DDB2* | FSFS | 93% |
| KM2 | *PPM1D DDB2 CCNF CDKN1A PCNA GADD45A PRKAB1 TOB1 TNFRSF10B MYC CCNB2 PTP4A1 BAX CCNA2 ATF3 LIG1 CCNG1 FHL2 PPP1R2 MBD4 RASGRP2 UBC NINJ1 TRIM22 IL2RB TP53BP1 PTPRCAP EEF1D PTPRE RAD23B EIF2B4 STX11 PTPN6 STK10 PSMD1 BTG3 MLH1 RNPEP HSPD1 UNG PTPRC PTPRA BCL2 GSS SH3BP5 TPP2 IDH3B CCNH STK11 EIF4EBP2 HSPA4 FADS2 RPA3 GZMK ANXA4 ICAM1 PPID LMO2 PPIE NUDT1 FUS POLR2A LY9 RPA1 PTS TNFRSF4 RPA2 PSMD8 GCDH MAN2C1 PTPN2 RUVBL1 ATP5H GK CD79B MAP4K4 POLE3 PRKCH AKT2 MOAP1 CCNG2 ALDOA SRD5A1 HAT1 XRCC1 EIF2S3 RAD1 UBE2A ZFP36L1 CD8A TALDO1 GPX4 SSBP2 ERCC3 ATP5O PEPD EIF4G2 ACO2 HEXB UBE3A ARPC1A PSMD10 PRCP PPIB ZNF337 CETN2 RPL29* | CSFS | 93% |
| **b) Derived from Radiation Dataset RadTBI-3 (GSE10640[GPL6522])** |  |  |  |
| M1 | *DDB2 HSPD1 MAP4K4 GTF3A PCNA MDH2* | FSFS | 86% |
| M2 | *DDB2 GTF3A TNFRSF10B* | FSFS | 80% |
| KM3 | *DDB2 RAD17 PSMD9 LY9 PPIH PCNA MDH2 MOAP1 TP53BP1 PPM1D ATP5G1 BCL2L2 ENO2 PTP4A1 PSMD8 LIG1 FDPS OGDH CCNG1 PSMD1* | BSFS | 95% |
| KM4 | *DDB2 HSPD1 ICAM1 PTP4A1 GTF3A LY9* | FSFS | 92% |
| KM5 | *RAD17 TNFRSF10B PSMD9 LY9 PPIH PCNA ZNF337 MDH2 TP53BP1 PPM1D ZFP36L1 ATP5G1 ALDOA BCL2L2 ENO2 GADD45A PTP4A1 PSMD8 LIG1 ATP5O FDPS OGDH PSMD1* | BSFS | 95% |
| **c) Derived from Radiation Dataset RadTBI-2 (GSE6874[GPL4782])** |  |  |  |
| M3 | *DDB2 CD8A TALDO1 PCNA EIF4G2 LCN2 CDKN1A PRKCH ENO1 PPM1D* | BSFS | 88% |
| M4 | *DDB2 CD8A TALDO1 PCNA LCN2 CDKN1A PRKCH ENO1 GTF3A IL2RB NINJ1 BAX TRIM22 PRKDC GADD45A MOAP1 ARPC1B LY9 LMO2 STX11 TPP2 CCNG1 GABARAP BCL2 GSS FTH1* | BSFS | 92% |
| KM6 | *DDB2 PRKDC PRKCH IGJ* | FSFS | 98% |
| KM7 | *DDB2 PRKDC TPP2 PTPRE GADD45A* | FSFS | 98% |
| <sup>1</sup> Performance metrics for these radiation gene signatures were previously reported in Zhao et al. (2018a) Tables 2 (k-fold validated signatures [KM1-KM7]) and 3 (traditionally validated signatures [M1-M4]); FS – Feature Selection metrics |  |  |  |

**Table 3.**
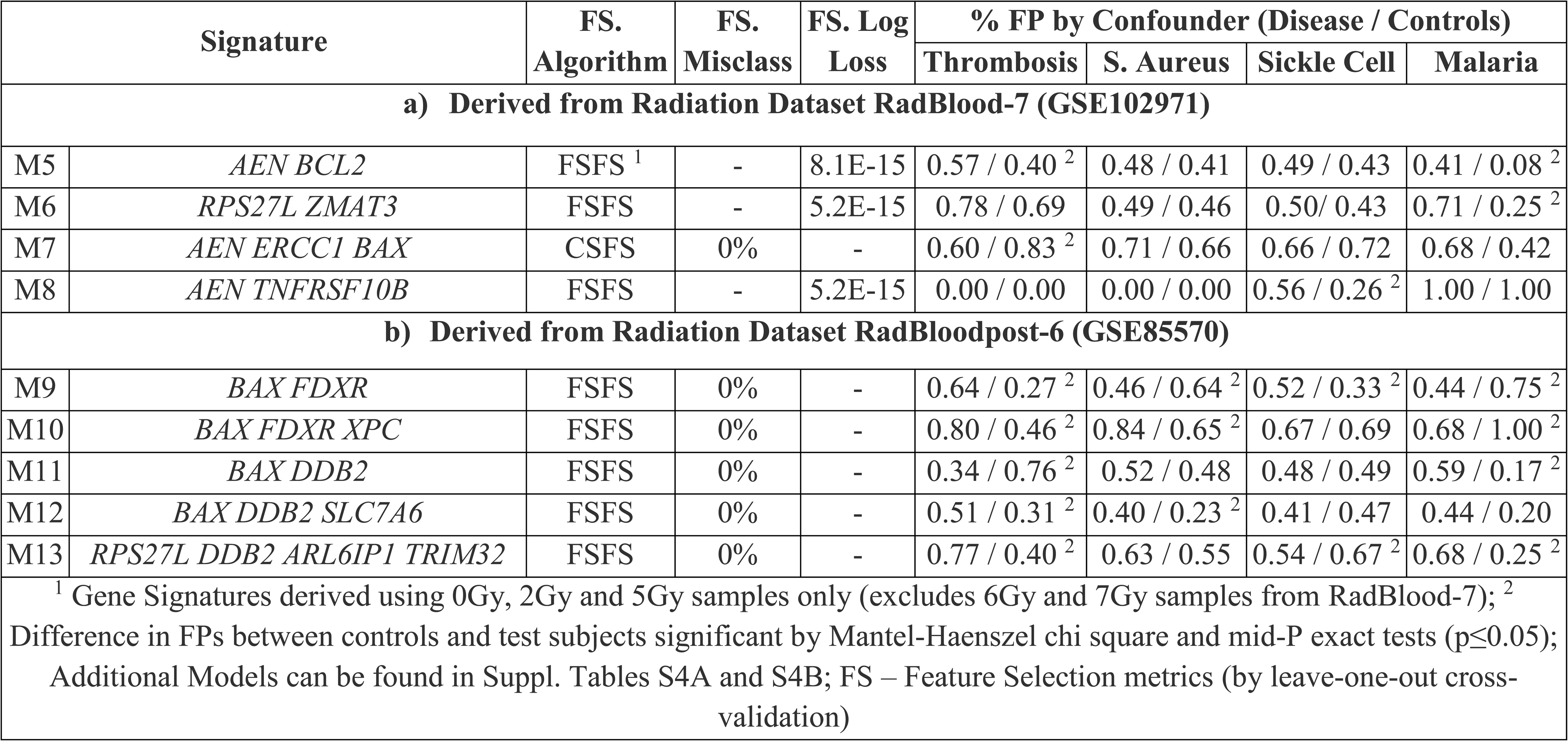
Radiation Gene Expression Signatures derived from RadBloodpost-6 and RadBlood-7 Radiation Datasets.

| Signature |  | FS. | FS. | FS. Log | % FP by Confounder (Disease / Controls) |  |  |  |
| --- | --- | --- | --- | --- | --- | --- | --- | --- |
|  |  | Algorithm | Misclass | Loss | Thrombosis | S. Aureus | Sickle Cell | Malaria |
| **a) Derived from Radiation Dataset RadBlood-7 (GSE102971)** |  |  |  |  |  |  |  |  |
| M5 | *AEN BCL2* | FSFS <sup>1</sup> | - | 8.1E-15 | 0.57 / 0.40 <sup>2</sup> | 0.48 / 0.41 | 0.49 / 0.43 | 0.41 / 0.08 <sup>2</sup> |
| M6 | *RPS27L ZMAT3* | FSFS | - | 5.2E-15 | 0.78 / 0.69 | 0.49 / 0.46 | 0.50 / 0.43 | 0.71 / 0.25 <sup>2</sup> |
| M7 | *AEN ERCC1 BAX* | CSFS | 0% | - | 0.60 / 0.83 <sup>2</sup> | 0.71 / 0.66 | 0.66 / 0.72 | 0.68 / 0.42 |
| M8 | *AEN TNFRSF10B* | FSFS | - | 5.2E-15 | 0.00 / 0.00 | 0.00 / 0.00 | 0.56 / 0.26 <sup>2</sup> | 1.00 / 1.00 |
| **b) Derived from Radiation Dataset RadBloodpost-6 (GSE85570)** |  |  |  |  |  |  |  |  |
| M9 | *BAX FDXR* | FSFS | 0% | - | 0.64 / 0.27 <sup>2</sup> | 0.46 / 0.64 <sup>2</sup> | 0.52 / 0.33 <sup>2</sup> | 0.44 / 0.75 <sup>2</sup> |
| M10 | *BAX FDXR XPC* | FSFS | 0% | - | 0.80 / 0.46 <sup>2</sup> | 0.84 / 0.65 <sup>2</sup> | 0.67 / 0.69 | 0.68 / 1.00 <sup>2</sup> |
| M11 | *BAX DDB2* | FSFS | 0% | - | 0.34 / 0.76 <sup>2</sup> | 0.52 / 0.48 | 0.48 / 0.49 | 0.59 / 0.17 <sup>2</sup> |
| M12 | *BAX DDB2 SLC7A6* | FSFS | 0% | - | 0.51 / 0.31 <sup>2</sup> | 0.40 / 0.23 <sup>2</sup> | 0.41 / 0.47 | 0.44 / 0.20 |
| M13 | *RPS27L DDB2 ARL6IP1 TRIM32* | FSFS | 0% | - | 0.77 / 0.40 <sup>2</sup> | 0.63 / 0.55 | 0.54 / 0.67 <sup>2</sup> | 0.68 / 0.25 <sup>2</sup> |
| <sup>1</sup> Gene Signatures derived using 0Gy, 2Gy and 5Gy samples only (excludes 6Gy and 7Gy samples from RadBlood-7); <sup>2</sup> Difference in FPs between controls and test subjects significant by Mantel-Haenszel chi square and mid-P exact tests (p≤0.05); Additional Models can be found in Suppl. Tables S4A and S4B; FS – Feature Selection metrics (by leave-one-out cross-validation) |  |  |  |  |  |  |  |  |

High levels of FP misclassification of viral infections were also evident with these signatures (Supp. Table S2A). KM6 and KM7 (derived from dataset RadTBI-2) misclassify all Influenza and most Dengue fever (BBD-Deng61, BBD-Deng08 and BBD-Deng78) datasets of patients at higher rates than uninfected controls. KM3-KM5 exhibited low FP rates in Influenza relative to other models, but Dengue virus datasets BBD-Deng62, BBD-Deng08 and BBD-Deng78 exhibited higher FP rates in infected samples relative to uninfected controls (Suppl. Table S2A). Interestingly, KM5 is the only gene signature in which *DDB2* is not present, and this gene contributes to high FP rates (Rogan et al. 2021). KM1 and KM2, which were derived from a third radiation dataset (RadLymphCL-1), often misclassified virus infected samples relative to controls (KM1 only: BBD-Deng61; KM2 only: BBD-Flu50, BBD-Flu31, BBD-Deng62 and BBD-Flu28; both KM1 and KM2: BBD-Deng08, BBD-Deng78, and BBD-Flu21). However, in some datasets, these models also demonstrated high FP rates in controls.

Expression levels in patients with latent Influenza A and Dengue fever stabilize at levels similar to uninfected controls after either convalescence or at the end stage infection. For example, M4 exhibited a 54% FP rate in Dengue-infected individuals 2-9 days after onset of symptoms (BBD-Deng08), but samples were correctly classified as unirradiated > 4 weeks after initial diagnosis (Figure 2A). The Influenza gene expression dataset BBD-Flu85 longitudinally sampled infected patients after initial symptoms at <72 hours [T1], 3-7 days [T2], and 2-5 weeks [T3]. FPs were significantly decreased at T3 for nearly all models tested (19 infected samples misclassified by M1 at T1 was reduced to 2 cases at T3; Figure 2B). These results clearly implicate these viral infections as the source of the transcriptional changes that affect parallel effects of radiation on these signature genes.

**Figure 2:**
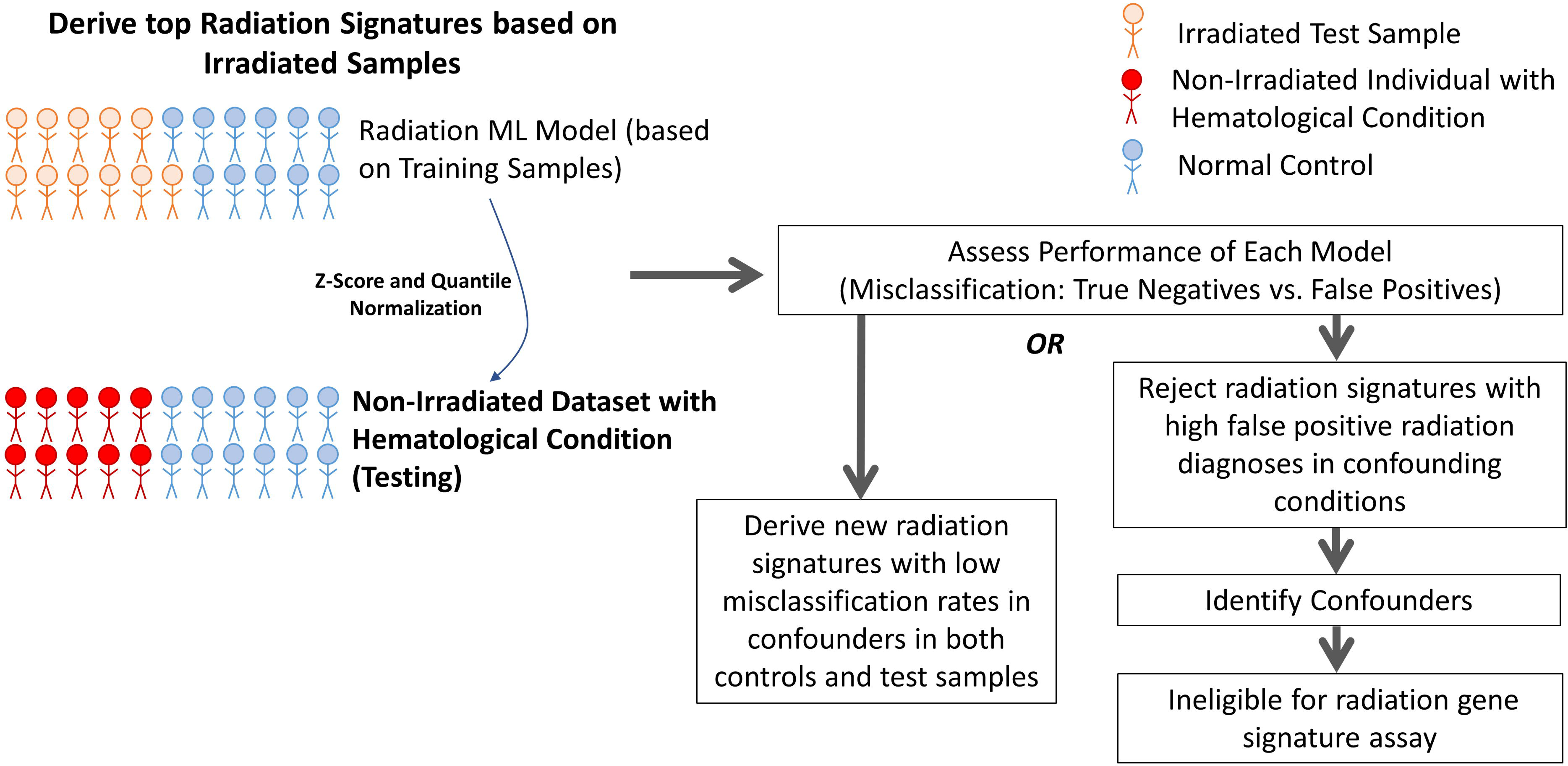
Performance of Traditionally-Validated Radiation Signatures on Confounders Stratified by Sub-phenotypes. Sankey diagrams delineate what fraction of disease patients and controls were properly (TN) and improperly classified (FP) by a radiation gene signature. A) The radiation signature M4 incorrectly classified 53% of Dengue-infected patient as irradiated, however all convalescent patients were properly classified (0% FP). B) Similarly, the FP rate of M1 decreased considerably after patient recovery (27% FP rate [N=19] against samples <3 days after symptoms; 3% [N=2] after 2-5 weeks). C) The FP rate of M3 was higher for severe malarial anemia patients versus those with cerebral malaria, suggesting that the differential expression caused by the two infection types may diverge in such a way that is measurable by M3. D) Conversely, the FP rate of venous thromboembolism patients by M4 was not influenced on whether the disease was recurrent.

### Specificity of radiation signatures using datasets of other hematological conditions

We investigated whether radiation gene signature accuracy was compromised by the presence of other blood borne infections and non-infectious, non-malignant hematological pathologies with publicly available expression data on patients with adequate sample sets (>10 individuals with corresponding control samples except for aplastic anemia). These included thromboembolism, S. aureus bacteremia, malaria, sickle cell disease, polycythemia vera, and aplastic anemia. We then determined recall levels for signatures M1-M4 and KM3-KM7 evaluated with these datasets, with the expectation that these models would predict all potential confounders as unirradiated (Suppl. Table S2B).

Each radiation gene signature was confounded by some, but not all, blood-borne disorders and infections. S. aureus infected samples were frequently misclassified as FPs by all signatures, except KM7 (Figure 3). High FP rates were observed for: M1 and KM5 – sickle cell and S. aureus; M2 – S. aureus; M3 and M4 – malaria, sickle cell, thromboembolism, polycythemia vera and S. aureus; KM3 and KM4 – malaria, sickle cell and S. aureus; KM6 – thromboembolism, polycythemia vera, S. aureus; KM7 – malaria, thromboembolism and polycythemia vera. We compared differences between FPs in patients and controls for each dataset using the Mantel- Haenszel chi square and mid-P exact statistical tests (Figure 3 and Suppl. Table S2B). Predictions of model M4 significantly confounded misclassification of radiation exposure for all conditions tested (polycythemia vera was only significant with the mid-P exact test), while the FP rate of KM6 was significantly higher in patients with either thrombosis or S. aureus infection (p-values indicated in Suppl. Table S2B). The malaria dataset stratified patients with either cerebral malaria or severe malarial anemia (Nallandhighal et al. 2019). The severe malarial anemia subset contains the majority of the FPs (Figure 2C). The thromboembolism dataset (Lewis et al. 2011), which categorized patient diagnoses as either single or recurrent thromboembolism, exhibited similar FP rates for both subsets (Figure 2D). Predictions by M3, M4, KM6 and KM7 were confounded by transcriptional changes resulting from different blood-borne conditions, while M2 and KM5 are the least influenced by these conditions. Aplastic anemia did not increase FP rates compared to controls for any of the signatures, consistent with our previous findings (Rogan et al. 2021).

**Figure 3:**
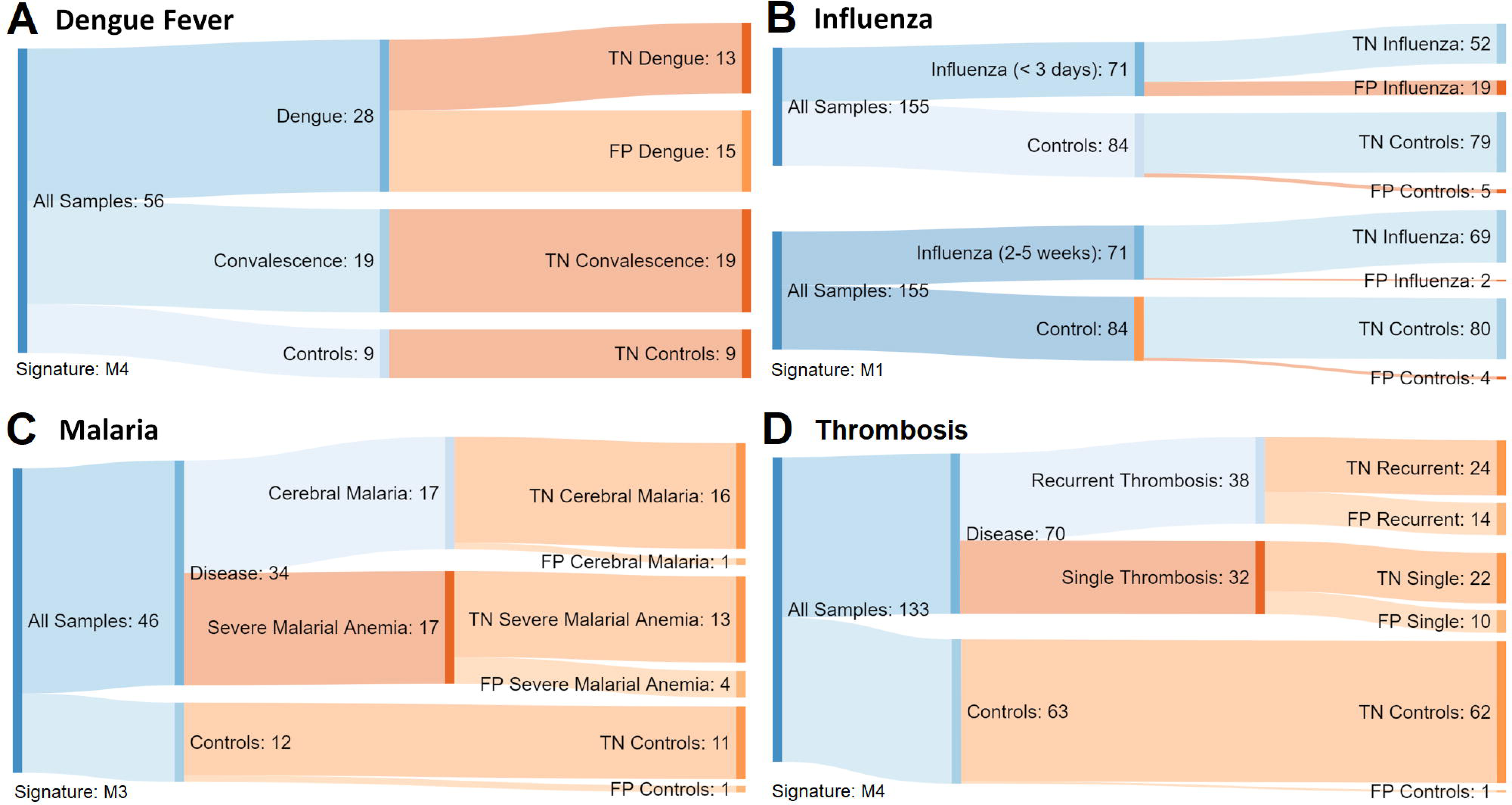
Misclassification Rates of Radiation Models in Unirradiated Blood-borne Disorders. Radiation gene signatures M1-M4 and KM3-KM7 performed well when predicting radiation exposure (≥80% overall accuracy; Table 2). However, many of these models falsely predicted individuals with blood-borne disorders (thromboembolism [A] and sickle cell disease [C]) and infectious diseases (S. aureus [B] and malaria [D]) as irradiated (%FP provided for individuals with the indicated disease [dark grey; top value], and controls [light grey; bottom value]). Asterisks indicate when the differences between the FP counts in controls and diseased individuals were significant with both Mantel-Haenszel chi square and mid-P exact tests (one-tailed; p<0.05). In general, the FP rate was high for all traditional validated (M1-M4) and most k-fold validated models (KM4, KM6 and KM7). Models KM3 and KM5 had a low FP rate across all datasets tested.

Predictions of radiation exposure by signatures M4, KM6 and KM7 were confounded by multiple viral, blood-borne infections and non-infectious blood disorders (Suppl. Table S2A and S2B). The genes responsible for the high sensitivity of these signatures were evident by comparative expression levels correctly (TN) vs incorrectly (FP) classified samples. The normalized gene expression distributions of TN and FP samples in malaria, S. aureus, sickle cell disease, thromboembolism, Influenza A and Dengue fever were visualized as violin plots (Figure 4 and Mucaki et al. 2021). The shared distributions in gene expression in FP confounders and radiation-exposed individuals can be observed without the need for advanced statistical measurements, however differences between TN and FP expression levels for the same gene were frequently also statistically significant. For example, expression of *BCL2* in sickle cell disease, and S. aureus and malaria infected samples was significantly lower in FP samples relative to TNs with M4 (p<0.05 with Student’s T-test, assuming two-tailed distribution and equal variance; Figure 4A [left]), similar to the effect of radiation exposure on expression of this gene (Figure 4A [right]). These same FP individuals have significantly higher *DDB2* expression in both S. aureus and sickle cell disease (Figure 4B). Increased *DDB2* expression was also observed for FPs using KM6 and KM7. For both genes, differences in expression in TN and FP samples were congruent with the changes observed in the radiation exposure datasets. Genes that may also contribute to misclassification include *GADD45A* in M4 (higher expression in diseased individuals vs. controls and induced by radiation exposure), and *PRKCH* and *PRKDC,* respectively, in KM6 and KM7 (decreased expression in FPs and in response to radiation). *BAX,* which is induced by radiation, is similarly expressed in FP and TN samples, and probably does not contribute to misclassification by M4.

**Figure 4:**
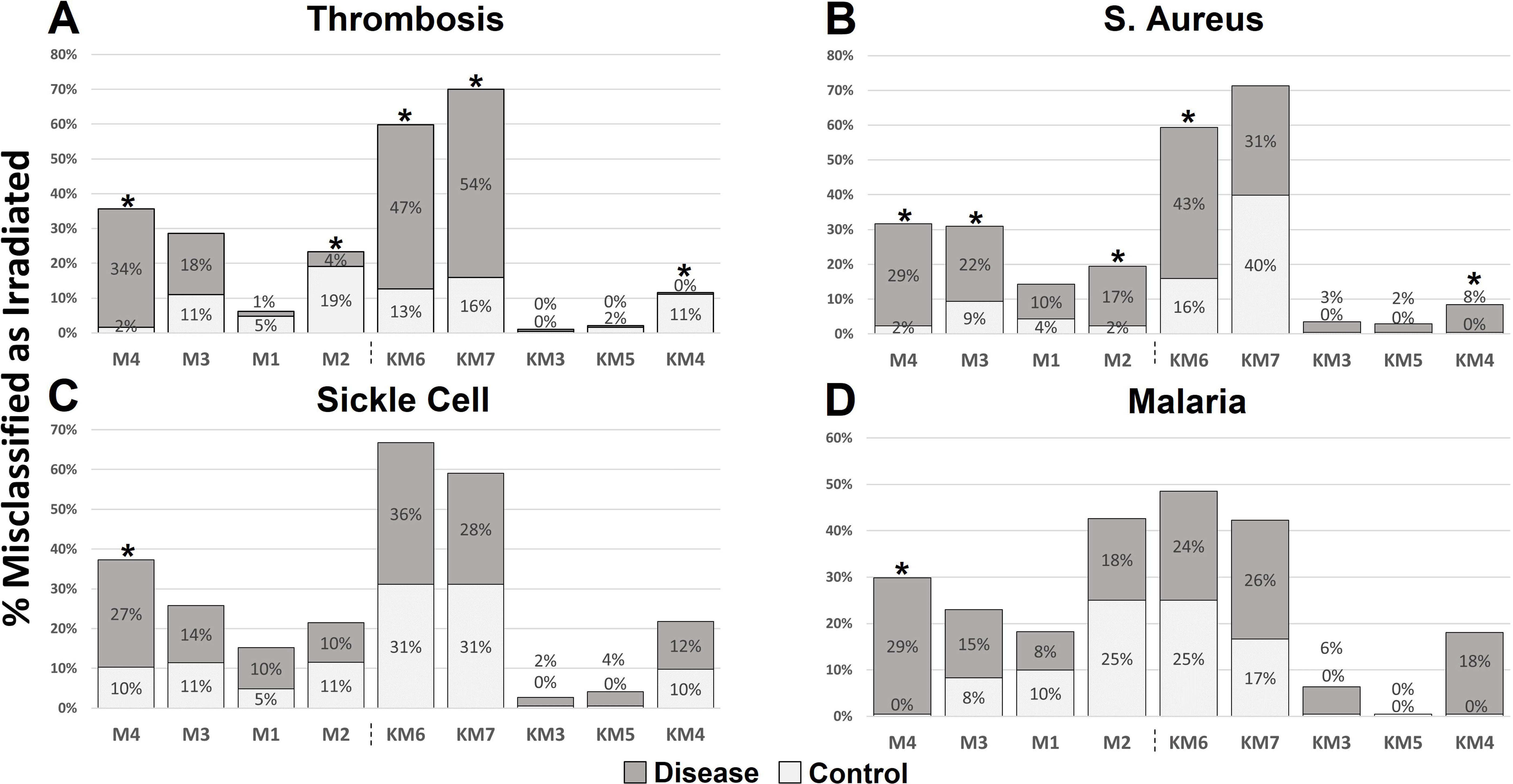
*DDB2* and *BCL2* Expression in Hematological Disorders with Radiation-Exposed Control Datasets. Normalized distribution of gene expression of confounder datasets (VTE: Venous Thromboembolism [orange]; SAu: S. aureus [teal]; Sic: Sickle Cell [yellow]; and Mal: Malaria [dark green]) for the genes **A)** *DDB2* and **B)** *BCL2* are presented as violin plots, where the expression of individuals with these conditions are divided by those predicted as irradiated (FP; left) or unirradiated (TN; right) by signature M4. Control expression of radiation-exposed (Irr.) and (Non.) unirradiated individuals are indicated by distributions labeled with light (dataset RadTBI-2) and dark (dataset RadTBI-3) red outlines on the right side of each panel. All expression differences between FP and TN samples (predicted with signature M4) found to be significant by Student’s t-test (assuming two-tailed distribution and equal variance) are indicated by brackets above the corresponding pair of predictions.

**Figure 5:**
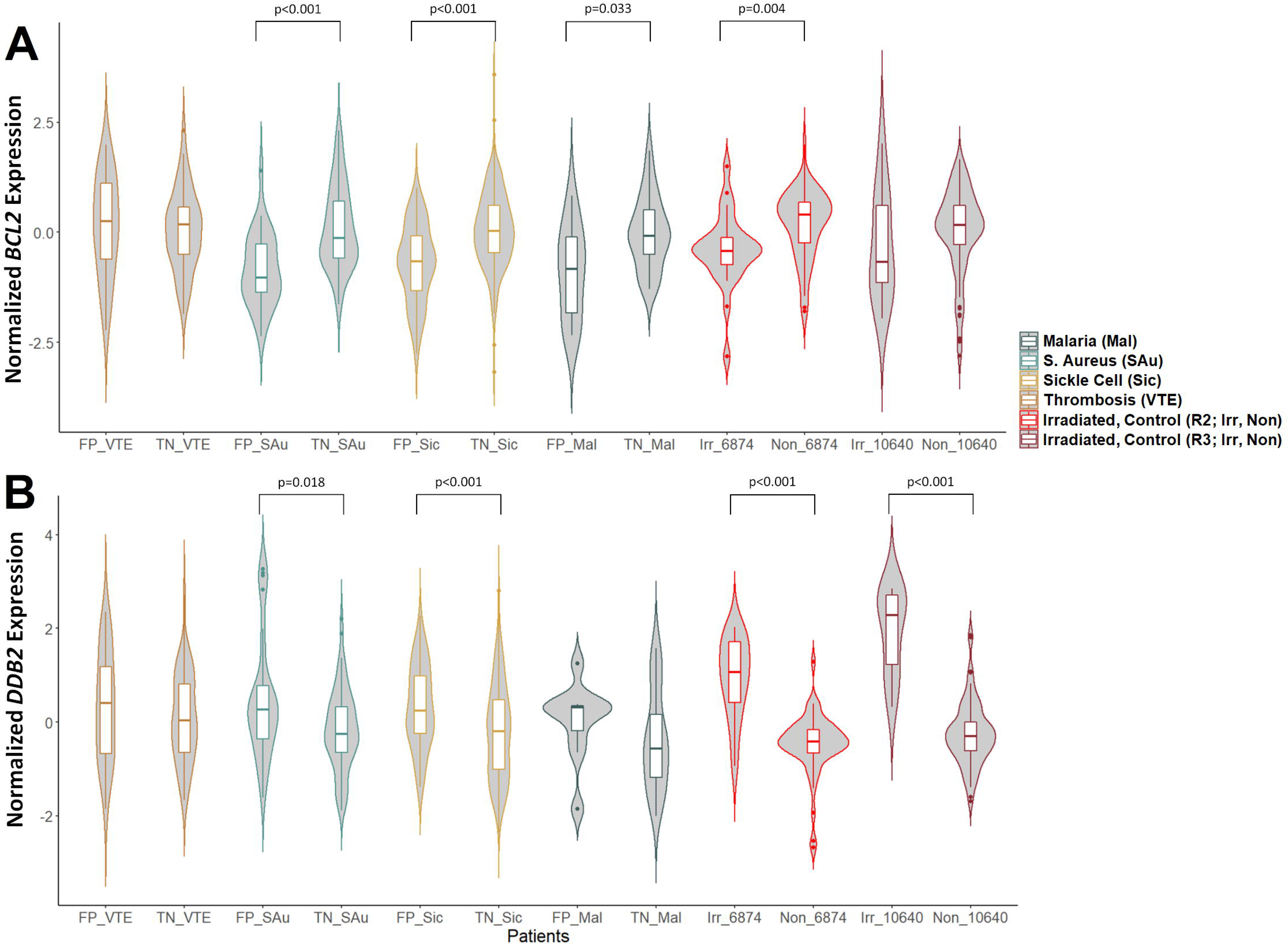
Multiple Genes Contribute to Misclassification of Confounding Datasets. Accuracy of M4 (Zhao et al. 2018a [Table 3B]) was significantly influenced by hematological confounders such as venous thromboembolism (top) and S. aureus infection (bottom). M4 misclassified diseased individuals (orange circles) far more often than controls (blue squares). Feature removal analysis of M4 determines if a particular gene was contributing to the %FP rate by observing how accuracy changes when a gene is removed. While M4 accuracy improved with the removal of *PRKDC*, *IL2RB* and *LCN2*, no individual gene restored misclassification back to control levels suggesting multiple genes are confounded by these diseases.

To determine the extent to which each gene contributes to the FP rates in each signature, gene features were removed individually, the radiation signature was rederived by biochemically-inspired ML, and misclassification rates were reassessed for each confounding condition (Suppl. Tables S3A [M1-M4] and S3B [KM3-KM7]). Removing any gene from gene signatures M1, M3, M4, KM3, KM5 and KM7 did not significantly alter the observed misclassification rates. Elimination of *PRKDC* (DNA double stranded break repair and recombination) and *IL2RB* (innate immunity/inflammation) reduced FP rates in thromboembolism patients by 10% and 5% for M4, respectively (Figure 5A), which still exceeded the FP rates of controls. Removal of these genes did not improve the FP rate of M4 in S. aureus-infected samples (Figure 5B). Thus, no single gene feature dominated the predictions by these signatures and could account for the misclassified samples. Removal of *DDB2*, *GTF3A* or *HSPD1* from KM4 significantly decreased its FP rate to the malaria dataset (18% to 0-3%; Suppl. Table S3B). Similarly, removal of *DDB2* from M2 and KM6 led to the complete elimination of FPs in both patients and controls. However, the removal of *DDB2* from these models was previously shown to severely reduce the true positive (TP) rate in irradiated samples (Zhao et al. 2018a); these genes cannot be eliminated without affecting the sensitivity of these signatures to accurately identify radiation exposed samples.

The contributions of individual signature genes can be assessed by evaluating their impact on overall model predictions for different patients. Expression changes were incrementally introduced to computationally determine the expression level required to change the outcome of the ML model (i.e. the inflection point of the prediction that distinguishes exposed from unirradiated samples). The threshold is visualized in the context of the expression value in the individual superimposed over a histogram of the distribution of expression for all confounders in the dataset. Individual expression values close to this threshold can indicate lower confidence in either the radiation exposure prediction or of misclassification by the model. Expression levels and thresholds of *DDB2*, *IL2RB*, *PCNA* and *PRKDC* for 3 individuals with thromboembolism (BBD- Thromb) predicted as irradiated by M4 (GSM474819, GSM474822, and GSM474828) are indicated in Suppl. Figure S1. Reduction of *DDB2* expression corrected misclassification for all patients, as did decreasing *PCNA* expression in GSM474822 and GSM474828. Increasing *IL2RB* and *PRKDC* expression of these two patients also corrected their misclassification. These results correspond to the effects of radiation on the expression of these genes in the RadTBI-2 and RadTBI-3 datasets, e.g. induction of *DDB2* and *PCNA*, repression of *IL2RB* and *PRKDC* (Mucaki et al. 2021). The expression changes required for *DDB2*, *PCNA* and *PRKDC* in these patients were nominal relative to the dynamic range of the entire dataset but were sufficient to alter predictions of the signatures. Conversely, changes in expression of *PCNA*, *IL2RB* or *PRKDC* were unable to modify the prediction of M4 in GSM474819. Only a large decrease in *DDB2* expression to levels below those of nearly all other thromboembolism patients was able to switch the classification of this individual. This reinforces previous observations about the strong impact of *DDB2* expression levels on prediction accuracy (Zhao et al. 2018a). Nevertheless, the combined expression of most of the genes which constitute the signature determine the classification result for each sample. Incorrect classifications where expression values are close to the model’s predictive inflection point are relevant when assessing misclassification accuracy. Generally, expression levels of most samples analyzed deviated significantly from these thresholds, leading to robust classifications by the M4 model.

### Misclassification of confounders with radiation gene signatures derived in this study

FSFS- and BSFS-based radiation gene signatures were derived from the top 50 ranked genes of datasets RadBloodpost-6 and RadBlood-7. RadBlood-7 contained sets of 20 samples, each irradiated at different absorbed energy levels (0 vs 2, 5, 6, and 7 Gy). Different ML models were derived either utilizing the full dataset or based on a combination of 2 and 5 Gy samples. The models derived from either subset of RadBlood-7 also exhibited very low misclassification (0-1 samples) and log-loss (<0.01). Common genes selected from signatures derived from RadBlood- 7 included *AEN*, *BAX*, *TNFRSF10B*, *RPS27L*, *ZMAT3* and *BCL2* (Table 3A and Suppl. Table S4A). Genes selected in RadBloodpost-6-based signatures included *BAX*, *FDXR*, *XPC*, *DDB2* and *TRIM32* (Table 3B and Suppl. Table S4B). All signatures from this dataset exhibited low misclassification rates (<0.5% by cross-validation).

The radiation gene signatures with the lowest misclassification rates from these datasets were evaluated against the blood-borne disease confounder datasets that compromised the accuracies of the M1-M4 and KM3-KM7 signatures (Zhao et al. 2018a). Misclassification rates were estimated using confounder datasets containing the largest numbers of samples, including thromboembolism, S. aureus infection, sickle cell disease and malaria. The signature designated M5 (consisting of *AEN* and *BCL2*; Table 3A) showed a significantly elevated FP rate over controls in blood samples from individuals with thromboembolism (18%) and malaria infection (33%; Suppl. Table S4A). Misclassification by M5 was increased by 6% in sickle cell disease, which also exhibited a significantly higher FP rate in an RadBlood-7-derived signature containing *AEN* (M8; Table 3A) in sickle cell disease (29%; p<0.05). Removal of genes from M8 significantly increased the FP rate for both controls and diseased individuals, which is a limitation of models based on small numbers of genes (Suppl. Table S5A). M9 (Table 3B) includes *BAX* and *FDXR,* and exhibited significantly increased FP rates in thromboembolism relative to controls (34-38% increased FP). Interestingly, M13 shows a significant increase in FPs of individuals with thromboembolism (similar to M1-M4), while M11 does not (Table 3B), despite both signatures containing *DDB2*. Removing any of the genes from these models did not substantially alter misclassification, except for a large decrease in FP upon removal of *RPS27L* from M9 (Suppl. Table S5B). Both M11 and M13 exhibited significantly high FP rates in malaria samples. BSFS models derived from dataset RadBloodpost-6 contained *FDXR*, *BAX* and *DDB2*, and showed high FP in S. aureus, sickle cell disease and malaria samples (significant by statistical analysis; Suppl. Table S4B). These confounders adversely affect the accuracy of gene signatures containing radiation response genes (such as *FDXR* and *AEN*) present in both these and other recently derived signatures in the published literature.

### Mitigating expression changes arising from confounding blood disorders with gene signatures comprising secreted factors originating in blood

Highly specific gene expression signatures that identify radiation exposed blood samples should also minimize inclusion of genes whose expression is altered by other hematological conditions. Predicted FPs in unexposed patients with confounding conditions may be the result of changes in expression of DNA damage and apoptotic genes that are shared with radiation responses. We derived ML-based gene signatures that exclude DNA damage or apoptotic genes which we anticipated would be less prone to misclassifying individuals with confounding blood disorders.

Changes in transcript levels of extracellular blood plasma proteins resulting from radiation exposure might exclude those associated with DNA damage or apoptosis response (for example, FLT3 ligand [*FLT3LG*] and amylase [AMY; *AMY1A, AMY2A*]; Barrett et al. 1982; Bertho et al. 2001; Tapio, 2013). This idea is predicated on observations that global protein synthesis significantly increases 4-8 hr after initial radiation exposures (Braunstein et al. 2009), with some profile changes detectable weeks to months later (Pernot et al. 2012; Hall et al. 2017). Radiation signatures in blood have been derived from proteins secreted into plasma (Wang et al. 2020) and expressed by multiple cell lineages (Ostheim et al. 2021). Radiation-induced short-term changes in the abundance of mRNAs encoding plasma proteins (that correspond to protein concentration changes) could allow steady state mRNA expression to be used as a surrogate for plasma protein levels. Significant correlations between mRNA and protein expression have been shown when the data have been transformed to normal distributions (Greenbaum et al. 2001; Greenbaum et al. 2002). This approach was adopted to derive mRNA signatures from radiation responsive genes in blood encoding secreted factors.

Genes which encode secreted proteins were used to derive new radiation gene expression signatures using our previously described methods (Zhao et al. 2018a). The plasma protein-encoding gene *GM2A* had the highest MI with radiation in dataset RadTBI-2 (MI=0.31), while *TRIM24* was highest in dataset RadTBI-3 (MI=0.27; Suppl. Table S1). *GM2A* is absent from dataset RadTBI-3. MI of *TRIM24* was low in RadTBI-2 (MI=0.05) resulting in it being ranked second to last (Suppl. Table S1) and it was not differentially expressed in this dataset (p-value > 0.05 by t-test; Suppl. Table S6C). Other top 50 ranked genes by mRMR in both datasets include *ACYP1*, *B4GALT5*, *FBXW7*, *IRAK3*, *MSRB2*, *NBL1*, *PRF1*, *SPOCK2*, and *TOR1A*.

We derived 5 independent radiation gene signatures encoding proteins secreted by blood cells (e.g. blood secretome models) that showed the lowest cross-validation misclassification accuracy or log-loss by various feature selection strategies (labeled SM1-SM5 [Secretome Model 1-5] in Table 4 and Suppl. Table S6A). SM5 feature selection was limited to the top 50 genes ranked by mRMR. This pre-selection step was not applied when deriving SM2 and SM3, whereas SM1 and SM4 were derived by CSFS feature selection which obtains genes sequentially by mRMR rank order without applying a threshold. Significantly upregulated genes and models consisted of *SLPI* (SM1, SM2), *TRIM24* (SM3, SM4, SM5), *TOR1A* (SM3, SM4), *GLA* (SM4), *SIL1* (SM4), *NUBPL* (SM4), *NME1* (SM4), *IPO9* (SM4), *IRAK3* (SM5), *MTX2* (SM5), and *FBXW7* (SM5). Downregulated genes included *CLCF1* (SM1, SM2), *USP3* (SM1, SM2), *TTC19* (SM2), *PFN1* (SM3, SM4, SM5), *CDC40* (SM4), *SPOCK2* (SM4), *CTSC* (SM4), *GLS* (SM4), and *PPP1CA* (SM5; Suppl. Table S6C). The models exhibited 12-39% misclassification (by k-fold validation) when validated against the alternative radiation dataset. The RadTBI-2 dataset was not suitable for signature derivation or validation, since data for *LCN2*, *ERP44*, *FN1*, *GLS*, and *HMCN1* were missing; these genes are present in models SM3 and/or SM4 (Table 4). The performance of the derived signatures was also assessed by inclusion of *FLT3* or *AMY*, either individually or in combination. These genes did not improve model accuracy beyond the levels of the best performing signatures that we derived.

**Table 4.**
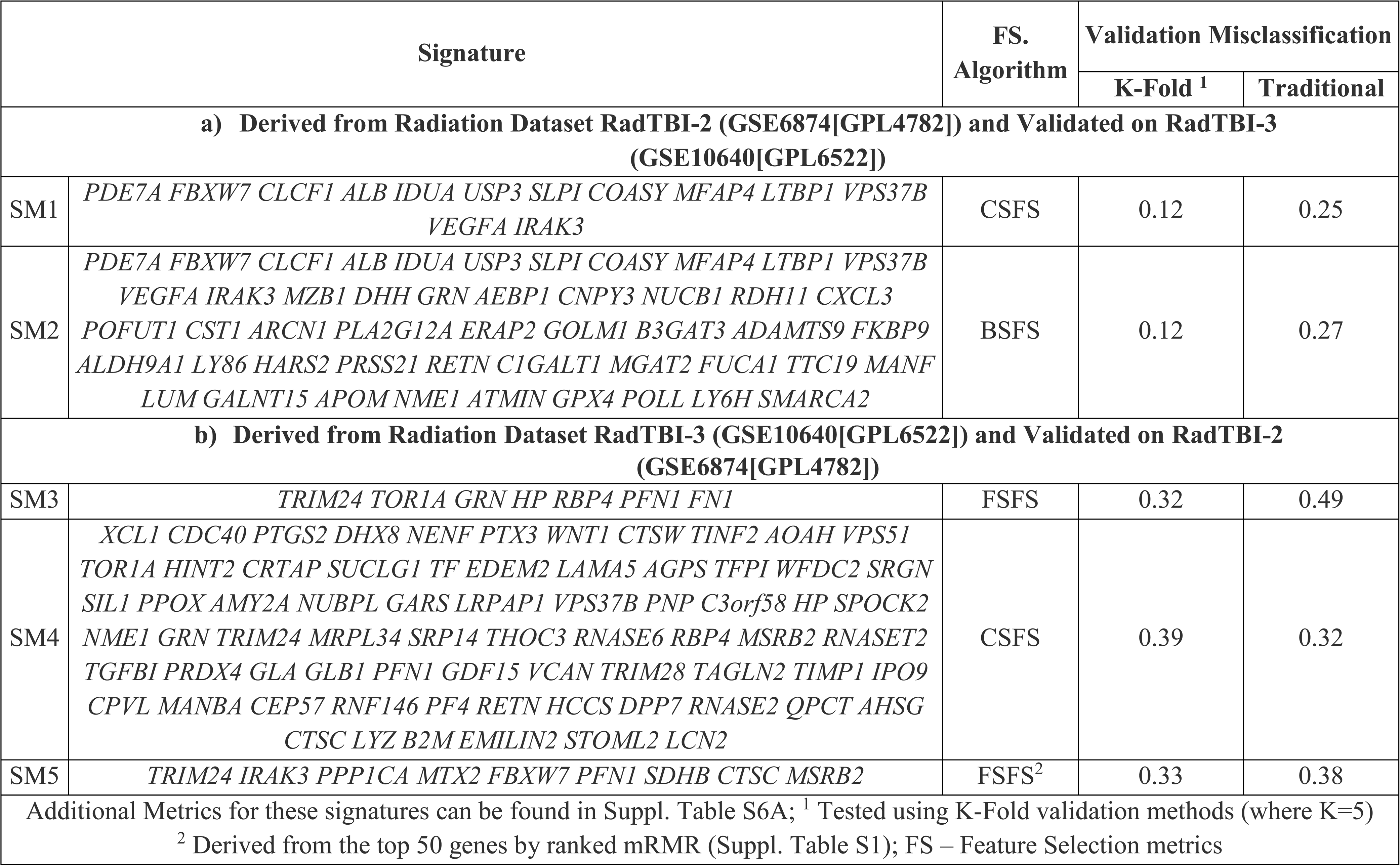
Radiation Gene Signatures including only Genes Encoding Secreted Factors derived from RadTBI-2 and RadTBI-3 Datasets.

The specificity of signatures derived from genes encoding secreted factors was then evaluated with expression data from unirradiated individuals with blood-borne diseases and infections (Suppl. Table S6B). SM3 and SM5 correctly classified nearly all samples in each dataset as unirradiated and maintained a FP rate <5% in all datasets (Figure 6). SM3 and SM5 contain <10 genes, were derived from dataset RadTBI-3 and share the genes *TRIM24* and *PFN1* (ranked #1 and #21 by mRMR). Both genes are significantly differentially expressed after radiation exposure, as is *TOR1A* in SM3 and *IRAK3*, *PPP1CA*, *MTX2*, *FBXW7* and *CTSC* in SM5 (Suppl. Table S6C). SM3 and SM5 have the highest fraction of genes found significant by Student’s t-test, which may explain its superior specificity relative to the other blood secretome signatures. Thromboembolism could only be evaluated with SM3 and SM5 due to missing genes from the SM1, SM2 and SM4 signatures. Conversely, SM1, SM2 and SM4 accuracy was compromised by expression changes of genes in one or more blood-borne diseases. Malaria (28%) and S. aureus (19%) infected patients were misclassified by SM1 as FPs (with 0.5% and 6.1% FP in controls, respectively), indicating that SM1 accuracy was significantly affected by these underlying infections (Suppl. Table S6B). SM4 accuracy was also impacted by S. aureus infection and sickle cell disease. Since the predictions of SM3 and SM5 were not influenced by the confounding conditions evaluated here, we suggest that these models will also be likely to be useful to exclude misclassification by others confounding conditions. Ultimately, it will be necessary to evaluate these signatures over a wide spectrum of other potential confounder phenotypes.

**Figure 6:**
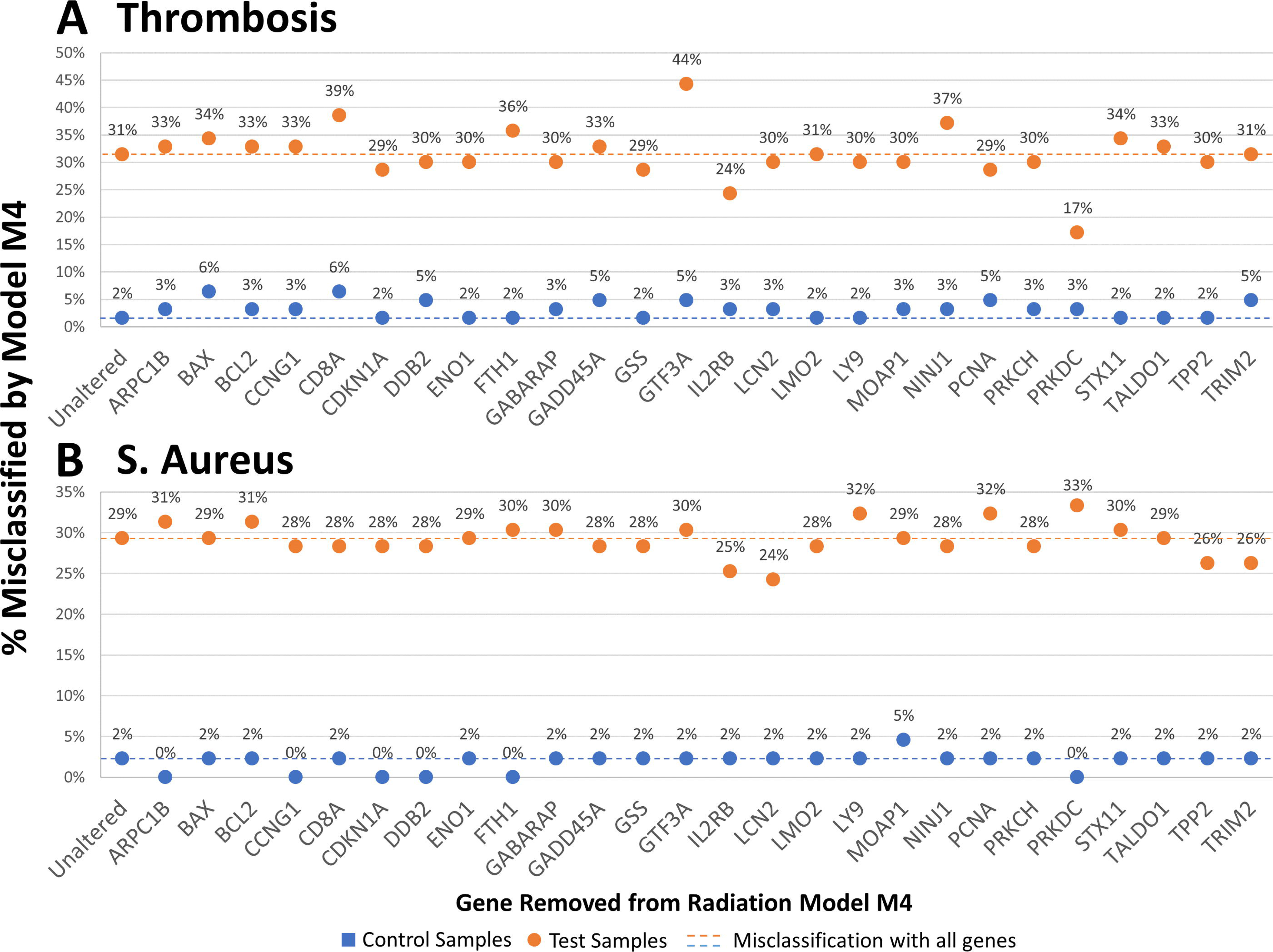
Radiation Gene Signatures derived from Transcripts Encoding Secreted Factors Reduce Misclassification in Unirradiated Confounder Phenotypes. Radiation signatures which consist exclusively of genes encoding for plasma secreted proteins were derived following the same basic approach of Zhao et al. (2018a). These models showed generally favorable performance when tested against an independent radiation dataset by k-fold validation (Table 4). Five blood secretome radiation signatures were derived consisting of 7-75 genes (SM1-SM5). Two models (SM3 and SM5) show high specificity across all hematological conditions tested (thromboembolism [A], S. aureus [B], sickle cell disease [C]) and malaria [D]).

SM3 and SM5 exhibited high specificity for radiation exposure (low false positivity in all confounding datasets) but were less sensitive than M1-M4 and KM3-KM7 (Table 4). Accurate identification of radiation exposed individuals should be feasible with a sequential strategy that first evaluates blood samples with suspected radiation exposures with signatures known to exhibit high sensitivity (e.g. M4; 88% accuracy to radiation exposure), followed by identification of FPs among predicted positives with SM3 and/or SM5 (which were not influenced by confounders; Suppl. Table S6B and S6D). By identifying and removing misclassified, unirradiated samples with the blood secretome-based radiation signatures, sequential application of both sets of signatures would predict predominantly TP samples.

## Discussion

We demonstrated high misclassification rates of radiation gene expression signatures in unirradiated individuals with either infections or blood borne disorders relative to normal controls. This was confirmed with a second set of k-fold validated radiation signatures from our previous study (Zhao et al. 2018a). Similar results were obtained with expression data from unirradiated individuals exhibiting other hematological conditions, which extended the spectrum of other abnormalities misclassified as exposed to radiation. Some of the same genes that are induced or repressed by radiation exhibit similar changes in direction and magnitude in infections and hematological conditions (for example, *DDB2*, *BCL2*). Signatures derived from more recent microarray platforms that contain key radiation response genes missing in our previous study (e.g. *FDXR, AEN*) were also prone to misclassifying hematological confounders as false positives. By assessing the performance of each model and rejecting signatures with a high rate of false radiation diagnoses in confounding conditions, many individuals with these comorbidities might be ineligible for these radiation gene signature assays. The symptoms of prodromal Influenza and ARS significantly overlap. During Influenza outbreaks, this could impact accurate and timely diagnosis of ARS. Expression-based bioassays might not improve this diagnostic accuracy, since traditional radiation signatures maximize sensitivity without accounting for the diminished specificity due to underlying hematological conditions. Other highly specific tests for radiation exposure, such as the dicentric chromosome assay, can be more accurate and less variable than expression-based assays, but require more time in the laboratory despite recent improvements in the speed of these analyses (Rogan et al. 2016; Liu et al. 2017; Shirley et al. 2017; Li et al. 2019, Shirley et al. 2020). Existing gene expression assays will need to address the false positive results obtained for individuals with hematological conditions before they can be used in general populations, who may not have a history of these conditions or who may have been pre-screened as a precondition to military or space deployment.

Use of matched, unirradiated controls provides a measure of sensitivity and dynamic range of the derived radiation gene signature. ML models for the same datasets can consist of different gene sets and are based on different C and sigma values, which can lead to differences in their ability to predict radiation exposures under different biological conditions. Nevertheless, genes with high mutual information between radiation amongst confounders consistently show differences in the distribution of TNs and FPs (Figure 4; Mucaki et al. 2021). That is, the models tend to unambiguously classify individual samples (Suppl. Fig. 1). Given the shared responses of different hematopathologies by leukocytes, the specificity of the signature for radiation exposure would, under ideal circumstances, be expected to exclude detection of other pathologies. Negative controls do not exhibit disease symptoms. In a nuclear incident or accident, the exposed population will include many individuals with underlying comorbidities. Application of radiation signatures derived by maximizing sensitivity in this population could lead to inappropriate diagnosis, and possibly treatment for ARS. The sequential gene signature assay design should improve the specificity of radiation gene expression assays in these individuals, and across the general population.

The cumulative incidences of these confounders are not rare, especially Influenza which affected approximately 11% of the US population during the 2019-2020 season (11,575 per 100,000; https://www.cdc.gov/flu/about/burden). The frequency of Dengue fever was also high in the Caribbean (2,510 per 100,000), Southeast Asia (2,940 per 100,000) and in South Asia (3,546 per 100,000; based on cases from 2017 [Zeng et al. 2017]). The annual prevalence of S. aureus bacteremia in the US is 38.2 to 45.7 per 100,000 person-years (El Atrouni et al. 2009; Rhee et al. 2015), but is higher among specific populations, such as hemodialysis patients. There are between 350,000 and 600,000 cases (200 per 100,000) of deep vein thromboembolism and pulmonary embolism that occur in the US every year (Anderson et al. 1991). Furthermore, there are over 100,000 individuals with sickle cell in the US (33.3 per 100,000; Hassell, 2010). Malaria is also common in sub-Saharan Africa in 2018 (21,910 per 100,000; World Health Organization, 2018). The prevalence of these diseases makes it clear that they could very well have a severe impact on assessment in a population-scale radiation exposure event.

Exploring the basis of these confounding disorders could facilitate strategies that minimize FPs suggesting radiation exposures. Common elements among their molecular etiologies may provide insight into their high misclassification rates. Despite their different clinical presentations, the underlying mechanisms of all of these conditions and radiation exposure appear to culminate in overwhelming chromosomal damage, degradation and cell lysis. Riboviral infections have been proposed to sequester host RNA binding proteins, leading to R-loop formation, DNA damage responses, and apoptosis (Rogan et al. 2021). This study suggested that expression of some key radiation signature genes appear to be altered by such infections. We also suggest that neutrophil extracellular traps (or NETs; Qi et al. 2020) may activate biochemical pathways that are present in early radiation responses. An early step in the formation of these structures is chromosome decondensation followed by the fragmentation of DNA which act as extracellular fibers which bind pathogens (such as S. aureus) in a process similar to autophagy in neutrophils (NETosis). This process would likely activate DNA damage in neutrophils, and some of the same DNA damage response genes that are activated (*DDB2*, *PCNA*, *GADD45A*) and repressed (*BCL2*) after radiation exposure are also similarly regulated after infections such as S. aureus. It has also been proposed that NET formation affects the severity of malaria infections (Boeltz et al., 2017). NETosis also contributes to the pathogenesis of numerous non-infectious diseases such as thromboembolism (Demers and Wagner, 2014; Collison, 2019) and sickle cell disease (Hounkpe et al. 2020), in addition to autoimmune disease (He et al. 2018) and general inflammation (Delgado-Rizo et al. 2017). If the origin of the FPs is confined to this lineage, then a comparison of the predictions of our traditionally validated signatures using data from the granulocyte versus lymphocyte lineages in individuals with these conditions should reveal whether NETosis is the likely etiology of the confounder expression phenotypes, or possibly even in radiation treated cells. To do this for radiation exposed cells, would require RNASeq data from these isolated cell populations (Ostheim et al. 2021). We would expect FPs in the confounder populations using signatures derived from myeloid-derived lineages, which include neutrophils.

The discovery of blood-borne conditions which lead to high FPs raises the question of whether other hematological conditions could also increase misclassification by radiation gene signatures. Such datasets are either unavailable or not suitable for analysis. Some studies consist with too few samples (e.g. GSE69601 [idiopathic portal hypertension] has 6 samples total) or lack the control samples necessary to perform a proper comparison (e.g. GSE33812 [aplastic anemia]). Although the available gene expression datasets covered a broad range of hematopathologies, additional testing of the sequential gene signatures will be required to exclude FPs due to underlying changes in gene expression from other confounders.

Confounding conditions will affect the precision of other assays and biomarkers that are routinely used to assess radiation exposure. Elevated levels of *γ*-H2AX, a marker of DNA damage, occur in cancer (Warters et al. 2005, Banath et al. 2004; Yu et al. 2006, Sedelnikova and Bonner 2006), in ulcerative colitis (Risques et al. 2008) and Zinc depletion/restriction (Mah et al. 2010). *γ*-H2AX has been suggested as an early cancer screening and cancer therapy biomarker (Sedelnikova and Bonner 2006). Besides its application for radiation assessment, the cytokinesis block micronucleus assay (CBMN; Fenech 2010) is also a multi-target endpoint for genotoxic stress from exogenous chemical agents (Kirsch-Volders et al. 2011; Fenech et al. 2016, Kirsch- Volders et al. 2018) and deficiency of micronutrients required for DNA synthesis and/or repair (folate, zinc; Beetstra et al. 2005; Sharif et al. 2012). The specificity of radiation testing may also be affected in patients with cancer using the *γ* -H2AX assay and patients under genotoxic stress and nutrient deficiencies using the CBMN assay.

Many radiation response genes were frequently selected as features for multiple signatures, and includes genes with roles in DNA damage response (*CDKN1A*, *DDB2*, *GADD45A*, *LIG1*, *PCNA*), apoptosis (*AEN*, *CCNG1*, *LY9*, *PPM1D, TNFRSF10B*), metabolism (*FDXR*), cell proliferation (*PTP4A1*) and the immune system (*LY9* and *TRIM22).* In general, the removal of these genes did not significantly alter the FP rate against confounder data. However, the removal of *LIG1*, *PCNA*, *PPM1D*, *PTP4A1*, *TNFRSF10B*, and *TRIM22* could partially decrease misclassification of Influenza samples in some models, as well as *DDB2* for Dengue (in addition to S. aureus and Polycythemia Vera). Many of these genes in our models are also present in other published radiation gene signatures and assays (Paul and Amundson, 2008; Lu et al. 2014; Oh et al. 2014; Port et al. 2017; Tichy et al. 2018; Jacobs et al. 2020). Paul and Amundsen (2008) developed a 74-gene radiation signature comprised of 16 genes present in the human signatures reported in Zhao et al. (2018a), including *CDKN1A*, *DDB2* and *PCNA*. Similarly, three of the 5 biomarkers implicated in Tichy et al. (2018) were also commonly selected (*CCNG1*, *CDKN1A*, and *GADD45A*), as were 5 of the 13 genes in the radiation assay described in Jacobs et al. (*BAX*, *CDKN1A*, *DDB2*, *MYC* and *PCNA*). While we cannot determine the impact on the accuracy of their signatures for confounders, it is evident that some genes that are included in these and other gene signatures (such as *DDB2*) can have a profound impact on the misclassification of individuals with confounding conditions.

The proposed sequential approach that combines highly sensitive predictors (affected by confounders) with high-specificity signatures could improve the accuracy of predicting TP exposures (Figure 7). After assessment with a sensitive signature (e.g. M4), all predicted positive samples would be re-evaluated with a high specificity signature (e.g. SM3 or SM5) to remove misclassified FP samples resulting in a higher performance assay that predominantly or exclusively labels truly irradiated samples. Datasets derived from different post-exposure times and exposures (RadLymphCL-1, RadTBI-2, RadTBI-3, RadBloodpost-6, and RadBlood-7) could generate ML models which can be used to assess the extent to which hematological confounders influence radiation exposure predictions for a variety of exposures and post-irradiation time constraints. A signature can be derived and selected which best fits the circumstances of the radiation exposure profile of a potentially exposed individual. The high specificity signatures, SM3 and SM5, were not as sensitive to misclassification as M1-M4 and KM1-KM7. It is conceivable that a proportion of radiation-exposed individuals could be misclassified as FNs if samples were evaluated solely with these signatures. Besides radiation exposure, the application of sequential gene signatures, each optimized respectively to maximize sensitivity and specificity, may turn out to be a general strategy for improving accuracy of molecular diagnoses for a wide spectrum of disease pathologies.

**Figure 7:**
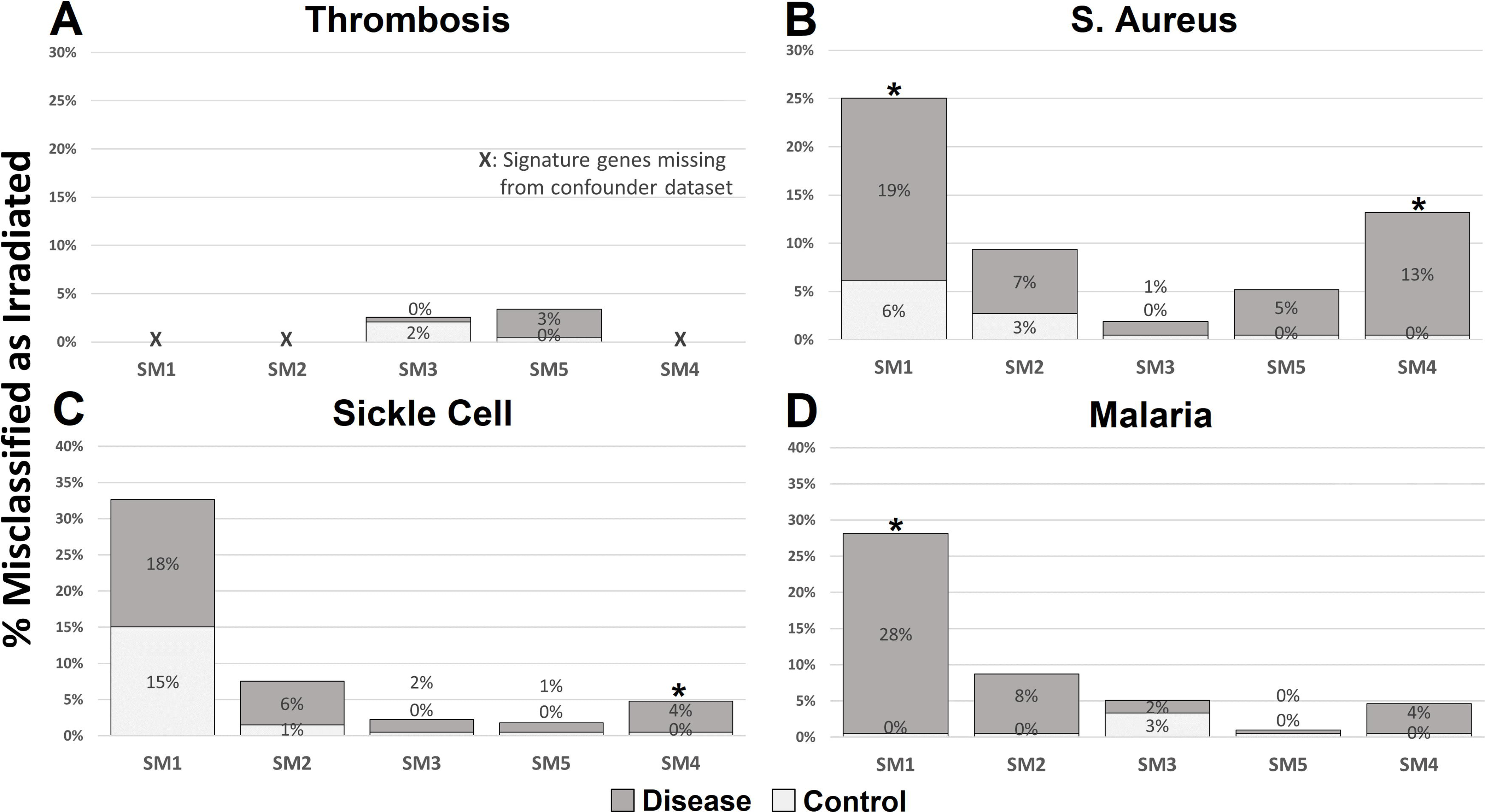
Sequential Application of Radiation-responsive and Blood Secretome Gene Signatures Identifies Exposed Individuals. False positive predictions due to differential expression caused by confounding conditions could be mitigated by following a sequential approach where samples are evaluated with both a highly sensitive radiation gene signature and a second signature with high specificity. M4, for example, is highly sensitive when validated against radiation dataset RadTBI-3 (88% accuracy), where all incorrect classifications were due to FP predictions (zero false negatives [FN]). Predicted irradiated samples could then be evaluated with a highly specific model such as SM3, which would identify and remove any misclassified unirradiated samples remaining in the set and leave only TPs.

Differential molecular diagnoses based on gene signatures would evaluate predicted radiation positive samples by individual gene signatures, each trained on different confounders (e.g. one model for Influenza infection, another for thromboembolism, etc). This approach would explicitly exclude FPs for radiation response while identifying the underlying condition. Separate signatures for sensitivity and specificity might also be avoided by training adversarial networks (Goodfellow et al. 2014) that contrast radiation-exposed samples with one or more datasets for confounding conditions, with emphasis on samples predicted to be FPs from the current radiation signatures. These resultant signatures would select radiation responsive genes which are resistant to the effects of confounders. Finally, ensuring that both the positive test and negative control samples in training sets properly account for the population frequencies of confounding diagnoses would also be expected to improve the performance of radiation gene signatures.

## Acknowledgements

This work was supported by the University of Western Ontario and CytoGnomix Inc. The authors thank Drs. Ruth Wilkins and Joan Knoll for their constructive comments.

## Data Availability Statement

A Zenodo data repository has been created for this study (DOI: <u>doi.org/10.5281/zenodo.5009008</u>). This archive provides additional violin plots which illustrate the expression of genes in models M1-M4, KM3-KM7 and SM1-SM5 for patients with a bloodborne condition or RNA viral infection.

---

**Supplemental Figure 1: Threshold Mapping of Venous Thromboembolism Patients Predicted Irradiated by Model M4**

The expression of *DDB2*, *PCNA*, *IL2RB* and *PRKDC* from all thrombosis patients (BBD-Thromb) (blue) and controls (orange) are presented as a histogram (non-normalized expression in 0.1 bins). The expression for patients GSM474822 (A), GSM474828 (B) and GSM474819 (C), each of which having been improperly classified as irradiated by M4, are indicated with a blue arrow. We iterated on patient gene expression for each gene and determined at which point the prediction switched from one class to the other (irradiated/unirradiated). The vertical dashed blue line designates the inflection point where expression of the gene will alter the prediction of the model. If the inflection point is absent, no change in gene expression could correct patient misclassification, which may indicate that the gene does not strongly contribute to the FP prediction of that individual.

## Supplementary Tables

Supplemental Table S1: mRMR Rankings of Radiation Response Genes in Ex-Vivo Control and Radiation Exposed Paired Datasets Supplemental Table S2A: Influenza and Dengue Infection Increase False Positives by Radiation Gene Signatures

Supplemental Table S2B: Infectious, Inherited and Non-Inherited Blood-borne Disorders Increase False Positives by Radiation Gene Signatures

Supplemental Table S3A: False Positive Rate after Feature Removal of M1-M4 Against Blood Disease Pathologies Supplemental Table S3B: False Positive Rate after Feature Removal of KM3-KM7 Against Blood Disease Pathologies

Supplemental Table S4A: Radiation Signatures derived from RadBlood-7 (GSE102971) with Increased FPs evaluating Blood Disease Pathologies

Supplemental Table S4B: Radiation Signatures derived from RadBloodpost-6 (GSE85570) with Increased FPs evaluating Blood Disease Pathologies

Supplemental Table S4C: Radiation Signatures derived from RadTBI-2 (GSE6874) with Increased FPs evaluating Blood Disease Pathologies

Supplemental Table S4D: Radiation Signatures derived from RadTBI-3 (GSE10640) with Increased FPs evaluating Diseased Blood Disease Pathologies

Supplemental Table S5A: Feature Removal Analysis Gene Signatures derived from Radiation Dataset RadBlood-7 (GSE102971) Supplemental Table S5B: Feature Removal Analysis Gene Signatures derived from Radiation Dataset RadBloodpost-6 (GSE85570) Supplemental Table S6A: Derivation of Radiation Models based on Genes Encoding Secreted Factors

Table S6B: Testing Blood Secretome Gene Radiation Signatures against Blood-Borne Diseased Patients

Supplementary Table S6C: Normalized change in mRNA expression of Blood Secretome gene signature components after Radiation Exposure

Supplemental Table S6D: Testing Blood Secretome Gene Radiation Signatures against Influenza and Dengue Infection

## References

Anderson FA Jr, Wheeler HB, Goldberg RJ, Hosmer DW, Patwardhan NA, Jovanovic B, Forcier A, Dalen JE. 1991. A population-based perspective of the hospital incidence and case-fatality rates of deep vein thrombosis and pulmonary embolism. The Worcester DVT Study. Arch Intern Med. 151(5):933–8.

Bagchee-Clark AJ, Mucaki EJ, Whitehead T, Rogan PK. 2020. Pathway-extended gene expression signatures integrate novel biomarkers that improve predictions of patient responses to kinase inhibitors. MedComm. 1:311–327.

Banáth JP, Macphail SH, Olive PL. 2004. Radiation sensitivity, H2AX phosphorylation, and kinetics of repair of DNA strand breaks in irradiated cervical cancer cell lines. Cancer Res. 64(19):7144–7149.

Banchereau R, Jordan-Villegas A, Ardura M, Mejias A, Baldwin N, Xu H, Saye E, Rossello-Urgell J, Nguyen P, Blankenship D, et al. 2012. Host immune transcriptional profiles reflect the variability in clinical disease manifestations in patients with Staphylococcus aureus infections. PLoS One. 7(4):e34390.

Barrett A, Jacobs A, Kohn J, Raymond J, Powles RL. 1982. Changes in serum amylase and its isoenzymes after whole body irradiation. Br Med J (Clin Res Ed) 285:170–171.

Beetstra S, Thomas P, Salisbury C, Turner J, Fenech M. 2005. Folic acid deficiency increases chromosomal instability, chromosome 21 aneuploidy and sensitivity to radiation-induced micronuclei. Mutat Res. 578(1-2):317–326.

Berdal JE, Mollnes TE, Wæhre T, Olstad OK, Halvorsen B, Ueland T, Laake JH, Furuseth MT, Maagaard A, Kjekshus H, et al. 2011. Excessive innate immune response and mutant D222G/N in severe A (H1N1) pandemic influenza. J Infect. 63(4):308–16.

Bertho JM, Demarquay C, Frick J, Joubert C, Arenales S, Jacquet N, Sorokine-Durm I, Chau Q, Lopez M, Aigueperse J, et al. 2001. Level of Flt3-ligand in plasma: a possible new bio-indicator for radiation-induced aplasia. Int J Radiat Biol. 77(6):703–12.

Boeltz S, Muñoz LE, Fuchs TA, Herrmann M. 2017. Neutrophil Extracellular Traps Open the Pandora’s Box in Severe Malaria. Front Immunol. 8:874.

Boldrini L, Bibault JE, Masciocchi C, Shen Y, Bittner MI. 2019. Deep Learning: A Review for the Radiation Oncologist. Front Oncol. 9:977.

Boldt S, Knops K, Kriehuber R, Wolkenhauer O. 2012. A frequency-based gene selection method to identify robust biomarkers for radiation dose prediction. Int J Radiat Biol. 88(3):267–76.

Braunstein S, Badura ML, Xi Q, Formenti SC, Schneider RJ. 2009. Regulation of Protein Synthesis by Ionizing Radiation. Mol. Cell. Biol. 29: 5645–56.

Budworth H, Snijders AM, Marchetti F, Mannion B, Bhatnagar S, Kwoh E, Tan Y, Wang SX, Blakely WF, Coleman M, et al. 2012. DNA repair and cell cycle biomarkers of radiation exposure and inflammation stress in human blood. PLoS One. 7(11):e48619.

Collison, J. 2019. Preventing NETosis to reduce thrombosis. Nat Rev Rheumatol. 15:317.

Cover TM, Thomas JA. 2006. Elements of Information Theory, 2nd edition. John Wiley & Sons, New York, NY, USA.

Cruz-Garcia L, O’Brien G, Donovan E, Gothard L, Boyle S, Laval A, Testard I, Ponge L, Woźniak G, Miszczyk L, et al. 2018. Influence of Confounding Factors on Radiation Dose Estimation Using In Vivo Validated Transcriptional Biomarkers. Health Phys. 115(1):90–101.

Delgado-Rizo V, Martínez-Guzmán MA, Iñiguez-Gutierrez L, García-Orozco A, Alvarado- Navarro A, Fafutis-Morris M. 2017. Neutrophil Extracellular Traps and Its Implications in Inflammation: An Overview. Front Immunol. 8:81.

Demers M, Wagner DD. 2014. NETosis: a new factor in tumor progression and cancer-associated thrombosis. Semin Thromb Hemost. 40(3):277–283.

Ding C, Peng H. 2005. Minimum redundancy feature selection from microarray gene expression data. J Bioinform Comput Biol. 3(2): 185–205.

Ding LH, Park S, Peyton M, Girard L, Xie Y, Minna JD, Story MD. 2013. Distinct transcriptome profiles identified in normal human bronchial epithelial cells after exposure to γ-rays and different elemental particles of high Z and energy. BMC Genomics. 14:372.

Disease Burden of Influenza. 2021. Centers for Disease Control and Prevention, National Center for Immunization and Respiratory Diseases (NCIRD). [accessed 2021 April 9]. https://www.cdc.gov/flu/about/burden

Dorman, SN, Baranova K, Knoll JHM, Urquhart BL, Mariani G, Carcangiu ML, Rogan PK. 2016. Genomic signatures for paclitaxel and gemcitabine resistance in breast cancer derived by machine learning. Molecular oncology, 10(1), 85–100.

Dressman HK, Muramoto GG, Chao NJ, Meadows S, Marshall D, Ginsburg GS, Nevins JR, Chute JP. 2007. Gene expression signatures that predict radiation exposure in mice and humans. PLoS Med. 4(4):e106.

El Atrouni WI, Knoll BM, Lahr BD, Eckel-Passow JE, Sia IG, Baddour LM. 2009. Temporal trends in the incidence of Staphylococcus aureus bacteremia in Olmsted County, Minnesota, 1998 to 2005: a population-based study. Clin Infect Dis.49(12):e130–8.

Fenech M. 2010. The lymphocyte cytokinesis-block micronucleus cytome assay and its application in radiation biodosimetry. Health Phys. 98(2):234–243.

Fenech M, Knasmueller S, Bolognesi C, Bonassi S, Holland N, Migliore L, Palitti F, Natarajan AT, Kirsch-Volders M. 2016. Molecular mechanisms by which in vivo exposure to exogenous chemical genotoxic agents can lead to micronucleus formation in lymphocytes in vivo and ex vivo in humans. Mutat Res. 770(Pt A):12–25.

Ghandhi SA, Smilenov LB, Elliston CD, Chowdhury M, Amundson SA. 2015. Radiation dose- rate effects on gene expression for human biodosimetry. BMC Med Genomics. 8:22.

Goodfellow IJ, Pouget-Abadie J, Mirza M, Xu B, Warde-Farley D, Ozair S, Courville A, Bengio Y. 2014. Generative adversarial nets. arxiv:1406.2661.

Greenbaum D, Luscombe NM, Jansen R, Qian J, Gerstein M. 2001. Interrelating different types of genomic data, from proteome to secretome: ’oming in on function. Genome Res. 11: 1463–1468.

Greenbaum D, Jansen R, Gerstein M. 2002. Analysis of mRNA expression and protein abundance data: an approach for the comparison of the enrichment of features in the cellular population of proteins and transcripts. Bioinformatics. 18: 585–596.

Guy JB, Bertoletti L, Magné N, Rancoule C, Mahé I, Font C, Sanz O, Martín-Antorán JM, Pace F, Vela JR, et al. 2017. Venous thromboembolism in radiation therapy cancer patients: Findings from the RIETE registry. Crit Rev Oncol Hematol. 113:83–89.

Hall J, Jeggo PA, West C, Gomolka M, Quintens R, Badie C, Laurent O, Aerts A, Anastasov N, Azimzadeh O, et al. 2017. Ionizing radiation biomarkers in epidemiological studies - An update. Mutat Res. 771:59–84.

Hassell KL. 2010. Population estimates of sickle cell disease in the U.S. Am J Prev Med. 38(4S):S512–S521.

He Y, Yang FY, Sun EW. 2018. Neutrophil Extracellular Traps in Autoimmune Diseases. Chin Med J (Engl). 131(13):1513–1519.

Hill A, Hanson M, Bogle MA, Duvic M. 2004. Severe radiation dermatitis is related to Staphylococcus aureus. Am J Clin Oncol. 27(4):361–363.

Hoang LT, Tolfvenstam T, Ooi EE, Khor CC, Naim AN, Ho EX, Ong SH, Wertheim HF, Fox A, Van Vinh Nguyen C, et al. 2014. Patient-based transcriptome-wide analysis identify interferon and ubiquination pathways as potential predictors of influenza A disease severity. PLoS One. 9(11):e111640.

Hounkpe BW, Chenou F, Domingos IF, Cardoso EC, Costa Sobreira MJV, Araujo AS, Lucena-Araújo AR, da Silva Neto PV, Malheiro A, Fraiji NA, et al. 2020. Neutrophil extracellular trap regulators in sickle cell disease: Modulation of gene expression of PADI4, neutrophil elastase, and myeloperoxidase during vaso-occlusive crisis. Res Pract Thromb Haemost. 16;5(1):204–210.

Jacobs AR, Guyon T, Headley V, Nair M, Ricketts W, Gray G, Wong JYC, Chao N, Terbrueggen R. 2020. Role of a high throughput biodosimetry test in treatment prioritization after a nuclear incident. Int J Radiat Biol. 96(1):57–66.

Jen KY, Cheung VG. 2003. Transcriptional response of lymphoblastoid cells to ionizing radiation. Genome Res. 13(9):2092–100.

Kirsch-Volders M, Plas G, Elhajouji A, Lukamowicz M, Gonzalez L, Vande Loock K, Decordier I. 2011. The in vitro MN assay in 2011: origin and fate, biological significance, protocols, high throughput methodologies and toxicological relevance. Arch Toxicol. 85(8):873–99.

Kirsch-Volders M, Fenech M, Bolognesi C. 2018. Validity of the Lymphocyte Cytokinesis-Block Micronucleus Assay (L-CBMN) as biomarker for human exposure to chemicals with different modes of action: A synthesis of systematic reviews. Mutat Res Genet Toxicol Environ Mutagen. 836(Pt A):47–52.

Knops K, Boldt S, Wolkenhauer O, Kriehuber R. 2012. Gene expression in low- and high-dose- irradiated human peripheral blood lymphocytes: possible applications for biodosimetry. Radiat Res. 178(4):304–12.

Kwissa M, Nakaya HI, Onlamoon N, Wrammert J, Villinger F, Perng GC, Yoksan S, Pattanapanyasat K, Chokephaibulkit K, Ahmed R, Pulendran B. 2014. Dengue virus infection induces expansion of a CD14(+)CD16(+) monocyte population that stimulates plasmablast differentiation. Cell Host Microbe. 16(1):115–27.

Lewis DA, Stashenko GJ, Akay OM, Price LI, Owzar K, Ginsburg GS, Chi JT, Ortel TL. 2011. Whole blood gene expression analyses in patients with single versus recurrent venous thromboembolism. Thromb Res. 128(6):536–40.

Li Y, Shirley BC, Wilkins RC, Norton F, Knoll JHM, Rogan PK. 2019. Radiation dose estimation by completely automated interpretation of the dicentric chromosome assay. Rad. Protect. Dosim. 186(1): 42–47.

Lipsitch M, Tchetgen Tchetgen E, Cohen T. 2010. Negative controls: a tool for detecting confounding and bias in observational studies [published correction appears in Epidemiology. 2010 Jul;21(4):589]. Epidemiology. 21(3):383–388.

Liu J, Li Y, Wilkins R, Flegal F, Knoll JHM, Rogan PK. 2017. Accurate cytogenetic biodosimetry through automated dicentric chromosome curation and metaphase cell selection [version 1; peer review: 2 approved]. F1000Res. 6:1396.

Lu TP, Hsu YY, Lai LC, Tsai MH, Chuang EY. 2014. Identification of gene expression biomarkers for predicting radiation exposure. Sci Rep. 4:6293.

Mah LJ, El-Osta A, Karagiannis TC. 2010. gammaH2AX: a sensitive molecular marker of DNA damage and repair. Leukemia. 24(4):679–686.

Meadows SK, Dressman HK, Muramoto GG, Himburg H, Salter A, Wei Z, Ginsburg GS, Chao NJ, Nevins JR, Chute JP. 2008. Gene expression signatures of radiation response are specific, durable and accurate in mice and humans. PLoS One. 3(4):e1912.

Mucaki EJ, Baranova K, Pham HQ, Rezaeian I, Angelov D, Ngom A, Rueda L, Rogan PK. 2016. Predicting Outcomes of Hormone and Chemotherapy in the Molecular Taxonomy of Breast Cancer International Consortium (METABRIC) Study by Biochemically-inspired Machine Learning [version 3; peer review: 2 approved]. F1000Research. 5:2124.

Mucaki EJ, Zhao J, Lizotte DJ, Rogan PK. 2019. Predicting responses to platin chemotherapy agents with biochemically-inspired machine learning. Signal transduction and targeted therapy. 4:1.

Mucaki EJ, Rogan PK. 2021. Zenodo Archive for " Improved radiation gene expression profiles with sequentially applied, sensitive and specific gene signatures". Zenodo. https://doi.org/10.5281/zenodo.5009008

Nallandhighal S, Park GS, Ho YY, Opoka RO, John CC, Tran TM. 2019. Whole-Blood Transcriptional Signatures Composed of Erythropoietic and NRF2-Regulated Genes Differ Between Cerebral Malaria and Severe Malarial Anemia. J Infect Dis. 219(1):154–164.

Oh DS, Cheang MC, Fan C, Perou CM. 2014. Radiation-induced gene signature predicts pathologic complete response to neoadjuvant chemotherapy in breast cancer patients. Radiat Res. 181(2):193–207.

Olagnier D, Peri S, Steel C, van Montfoort N, Chiang C, Beljanski V, Slifker M, He Z, Nichols CN, Lin R, et al. 2014. Cellular oxidative stress response controls the antiviral and apoptotic programs in dengue virus-infected dendritic cells. PLoS Pathog. 10(12):e1004566.

Ostheim P, Don Mallawaratchy A, Müller T, Schüle S, Hermann C, Popp T, Eder S, Combs SE, Port M, Abend M. 2021. Acute radiation syndrome-related gene expression in irradiated peripheral blood cell populations. Int J Radiat Biol. 97(4):474–484.

Park JG, Paul S, Briones N, Zeng J, Gillis K, Wallstrom G, LaBaer J, Amundson SA. 2017. Developing Human Radiation Biodosimetry Models: Testing Cross-Species Conversion Approaches Using an Ex Vivo Model System. Radiat Res. 187(6):708–721.

Paul S, Amundson SA. 2008. Development of gene expression signatures for practical radiation biodosimetry. Int J Radiat Oncol Biol Phys. 71(4):1236–1244.

Paul S, Amundson SA. 2011. Gene expression signatures of radiation exposure in peripheral white blood cells of smokers and non-smokers. Int J Radiat Biol. 87(8):791–801.

Pernot E, Hall J, Baatout S, Benotmane MA, Blanchardon E, Bouffler S, El Saghire H, Gomolka M, Guertler A, Harms-Ringdahl M, et al. 2012. Ionizing radiation biomarkers for potential use in epidemiological studies. Mutat Res. 751(2):258–286.

Port M, Hérodin F, Valente M, Drouet M, Lamkowski A, Majewski M, Abend M. 2017. Gene expression signature for early prediction of late occurring pancytopenia in irradiated baboons. Ann Hematol. 96(5):859–870.

Qi J-L, He J-R, Liu C-B, Jin S-M, Gao R-Y, Yang X, Bai H-M, Ma Y-B. 2020. Pulmonary Staphylococcus aureus infection regulates breast cancer cell metastasis via neutrophil extracellular traps (NETs) formation. MedComm. 1:188–201.

Quinlan J, Idaghdour Y, Goulet JP, Gbeha E, de Malliard T, Bruat V, Grenier JC, Gomez S, Sanni A, Rahimy MC, Awadalla P. 2014. Genomic architecture of sickle cell disease in West African children. Front Genet. 5:26.

Rhee Y, Aroutcheva A, Hota B, Weinstein RA, Popovich KJ. 2015. Evolving Epidemiology of Staphylococcus aureus Bacteremia. Infect Control Hosp Epidemiol. 36(12):1417–22.

Rieger KE, Hong WJ, Tusher VG, Tang J, Tibshirani R, Chu G. 2004. Toxicity from radiation therapy associated with abnormal transcriptional responses to DNA damage. Proc Natl Acad Sci U S A. 101(17):6635–40.

Risques RA, Lai LA, Brentnall TA, Li L, Feng Z, Gallaher J, Mandelson MT, Potter JD, Bronner MP, Rabinovitch PS. 2008. Ulcerative colitis is a disease of accelerated colon aging: evidence from telomere attrition and DNA damage. Gastroenterology. 135(2):410–8.

Rogan PK, Li Y, Wilkins RC, Flegal FN, Knoll JH. 2016. Radiation Dose Estimation by Automated Cytogenetic Biodosimetry. Radiat Prot Dosimetry. 172(1-3):207–217.

Rogan PK. 2019. Multigene signatures of responses to chemotherapy derived by biochemically- inspired machine learning. Mol Genet Metab. 128(1-2):45–52.

Rogan PK, Mucaki EJ, Shirley BC. 2020. Characteristics of human and viral RNA binding sites and site clusters recognized by SRSF1 and RNPS1. Zenodo. http://www.doi.org/10.5281/zenodo.3737089

Rogan PK, Mucaki EJ and Shirley BC. 2021. A proposed molecular mechanism for pathogenesis of severe RNA-viral pulmonary infections [version 2; peer review: 4 approved]. F1000Research. 9:943.

Sedelnikova OA, Bonner WM. 2006. GammaH2AX in cancer cells: a potential biomarker for cancer diagnostics, prediction and recurrence. Cell Cycle. 5:2909–2913.

Sharif R, Thomas P, Zalewski P, Fenech M. 2012. Zinc deficiency or excess within the physiological range increases genome instability and cytotoxicity, respectively, in human oral keratinocyte cells. Genes Nutr. 7(2):139–154.

Shirley B, Li Y, Knoll JHM, Rogan PK. 2017. Expedited Radiation Biodosimetry by Automated Dicentric Chromosome Identification (ADCI) and Dose Estimation. J Vis Exp. (127):56245.

Shirley BC, Knoll JHM, Moquet J, Ainsbury E, Pham ND, Norton F, Wilkins RC, Rogan PK. 2020. Estimating partial-body ionizing radiation exposure by automated cytogenetic biodosimetry. Int J Radiat Biol. 96(11):1492–1503.

Smirnov DA, Brady L, Halasa K, Morley M, Solomon S, Cheung VG. 2012. Genetic variation in radiation-induced cell death. Genome Res. 22(2):332–9.

Spivak JL, Considine M, Williams DM, Talbot CC Jr, Rogers O, Moliterno AR, Jie C, Ochs MF. 2014. Two clinical phenotypes in polycythemia vera. N Engl J Med. 371(9):808–17.

Svensson JP, Stalpers LJ, Esveldt-van Lange RE, Franken NA, Haveman J, Klein B, Turesson I, Vrieling H, Giphart-Gassler M. 2006. Analysis of gene expression using gene sets discriminates cancer patients with and without late radiation toxicity. PLoS Med. 3(10):e422.

Tang BM, Shojaei M, Parnell GP, Huang S, Nalos M, Teoh S, O’Connor K, Schibeci S, Phu AL, Kumar A, et al. 2017. A novel immune biomarker *IFI27* discriminates between influenza and bacteria in patients with suspected respiratory infection. Eur Respir J. 49(6):1602098.

Tapio S. 2013. Ionizing Radiation Effects on Cells, Organelles and Tissues on Proteome Level. pp 37–48 In: Leszczynski D. (eds) Radiation Proteomics. Advances in Experimental Medicine and Biology, vol 990. Springer, Dordrecht.

Tian Y, Babor M, Lane J, Schulten V, Patil VS, Seumois G, Rosales SL, Fu Z, Picarda G, Burel J, et al. 2017. Unique phenotypes and clonal expansions of human CD4 effector memory T cells re- expressing CD45RA. Nat Commun. 8(1):1473.

Tichy A, Kabacik S, O’Brien G, Pejchal J, Sinkorova Z, Kmochova A, Sirak I, Malkova A, Beltran CG, Gonzalez JR, et al. 2018. The first in vivo multiparametric comparison of different radiation exposure biomarkers in human blood. PLoS One. 13(2):e0193412.

Tsuge M, Oka T, Yamashita N, Saito Y, Fujii Y, Nagaoka Y, Yashiro M, Tsukahara H, Morishima T. 2014. Gene expression analysis in children with complex seizures due to influenza A(H1N1)pdm09 or rotavirus gastroenteritis. J Neurovirol. 20(1):73–84.

van Oorschot B, Uitterhoeve L, Oomen I, Ten Cate R, Medema JP, Vrieling H, Stalpers LJ, Moerland PD, Franken NA. 2017. Prostate Cancer Patients with Late Radiation Toxicity Exhibit Reduced Expression of Genes Involved in DNA Double-Strand Break Repair and Homologous Recombination. Cancer Res. 77(6):1485–1491.

Vanderwerf SM, Svahn J, Olson S, Rathbun RK, Harrington C, Yates J, Keeble W, Anderson DC, Anur P, Pereira NF, et al. 2009. TLR8-dependent TNF-(alpha) overexpression in Fanconi anemia group C cells. Blood. 114(26):5290–8.

Wang Q, Lee Y, Shuryak I, Pujol Canadell M, Taveras M, Perrier JR, Bacon BA, Rodrigues MA, Kowalski R, Capaccio C, et al. 2020. Development of the FAST-DOSE assay system for high- throughput biodosimetry and radiation triage. Sci Rep. 10(1):12716.

Warters RL, Adamson PJ, Pond CD, Leachman SA. 2005. Melanoma cells express elevated levels of phosphorylated histone H2AX foci. J Invest Dermatol. 124:807–817.

Yu T, MacPhail SH, Banath JP, Klokov D, Olive PL. 2006. Endogenous expression of phosphorylated histone H2AX in tumors in relation to DNA double-strand breaks and genomic instability. DNA Repair (Amst). 5:935–946.

Zeng G. 2015. A Unified Definition of Mutual Information with Applications in Machine Learning. Math. Probl. Eng. 2015, 1–12.

Zeng Z, Zhan J, Chen L, Chen H, Cheng S. 2021. Global, regional, and national dengue burden from 1990 to 2017: A systematic analysis based on the global burden of disease study 2017. E Clinical Medicine. 32:100712.

Zhao JZL, Mucaki EJ Rogan PK. 2018a. Predicting ionizing radiation exposure using biochemically-inspired genomic machine learning [version 2; peer review: 3 approved]. F1000Research. 7:233.

Zhao JZL, Mucaki EJ, Rogan PK. 2018b. Matlab Code for “Predicting Exposure to Ionizing Radiation by Biochemically-Inspired Genomic Machine Learning”. Zenodo. https://doi.org/10.5281/zenodo.1170571

