## Supplementary figures and images for "Improved radiation expression profiling in blood by sequential application of sensitive and specific gene signatures"

### Supplemental Figure 1

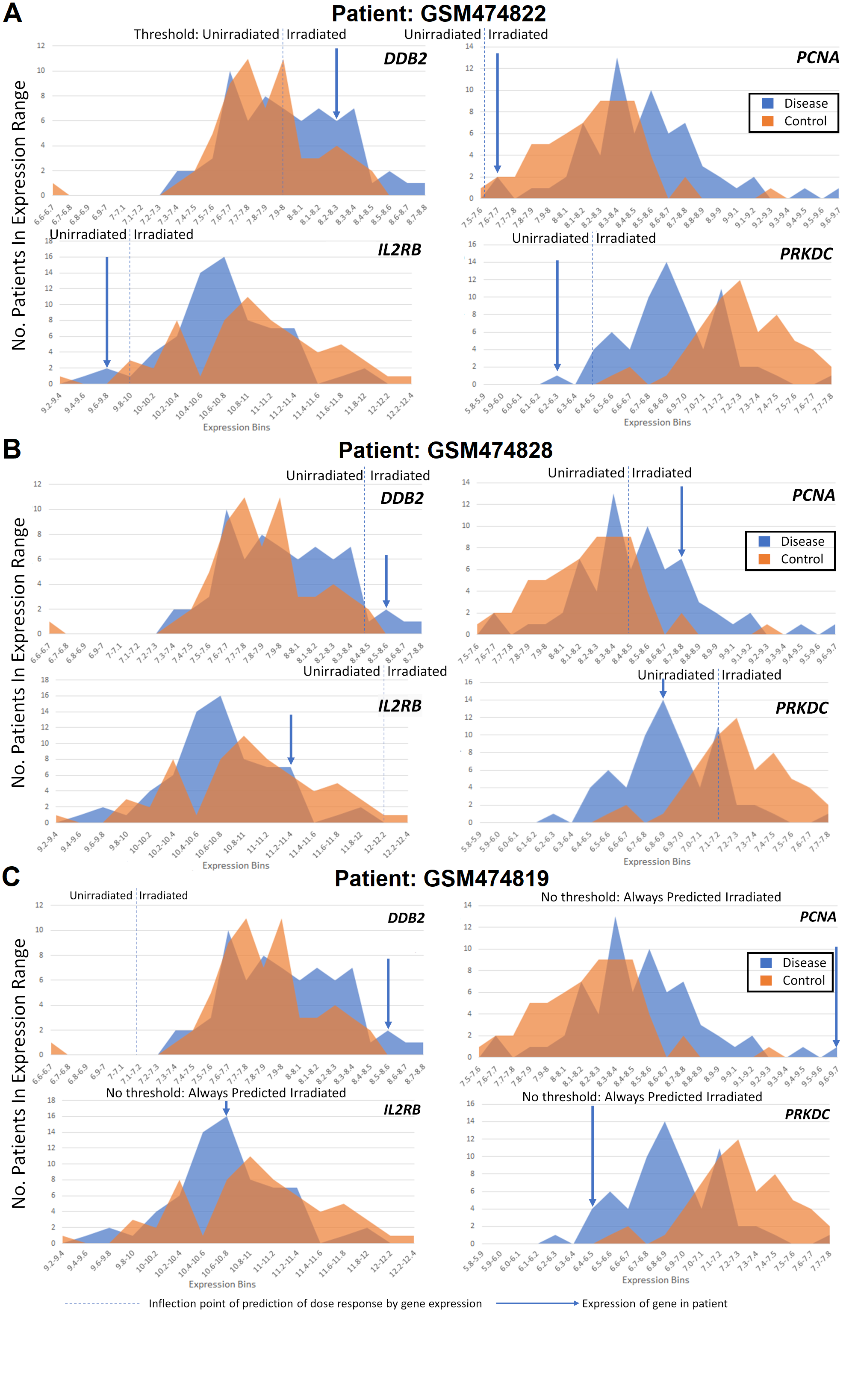
